# Selective depletion of upper-layer somatostatin interneuron subtypes in schizophrenia

**DOI:** 10.64898/2026.09.18.752621

**Authors:** Nicole Endresz, Maria Eleni Fafouti, Keon Arbabi, Xiaolin Zhou, Thomas DeLong, Guillermo Gonzalez-Burgos, Laramie Duncan, Etienne Sibille, Shreejoy J Tripathy

## Abstract

Schizophrenia (SCZ) is associated with cortical GABAergic dysfunction, but whether inhibitory interneurons are lost or persist in an altered molecular state remains unresolved. Here, we harmonized seven prefrontal post-mortem single-nucleus RNA-seq datasets onto a fine-grained taxonomy of cortical cell types and meta-analyzed their gene expression and abundance changes in SCZ (298 controls, 171 SCZ). First, we find a subclass-wide reduction of *SST* mRNA within somatostatin (Sst) neurons. Second, we find reduced abundance (depletion) of a subset of upper-layer Sst interneurons and increased abundance of L6b excitatory neurons, with both changes confirmed in spatial transcriptomics (12 controls, 12 SCZ). Notably, SCZ genetic risk is enriched in the most depleted Sst cells. Depleted Sst subtypes highly express *HCN1*, partially correspond to primate-specialized *CALB1*-expressing double-bouquet cells, and are among the cells lost earliest in Alzheimer’s disease. These upper-layer Sst interneurons constitute an intrinsically vulnerable population and a promising target for neuroprotective and compensatory therapies.

## Introduction

Schizophrenia (SCZ) is a chronic neuropsychiatric disorder affecting approximately 0.5-1% of adults worldwide (Saha et al. 2005). It is characterized by positive symptoms such as hallucinations and delusions, negative symptoms such as blunted affect and anhedonia, and pervasive cognitive impairment (Kahn et al. 2015). These symptoms contribute to difficulties in social integration and to disparities in health-related outcomes, including markedly reduced life expectancy (Hjorthøj et al. 2017). SCZ typically requires lifelong treatment and available drug therapies are principally second-generation antipsychotics that target dopaminergic signaling; however, approximately 10-30% of SCZ patients display treatment resistance, and a further 30-60% experience only partial improvement or intolerable side effects, pointing to the need for alternative drug targets (Siskind et al. 2022; Howes et al. 2017). Moreover, antipsychotic therapy does not improve, and might indeed affect, cognitive function in SCZ (Feber et al. 2025). Progress toward novel therapies has been slowed by an incomplete picture of the specific cortical cell types and molecular changes that underlie the disorder (Hyman 2012).

Unlike Alzheimer’s or Parkinson’s disease, SCZ does not involve gross neurodegeneration (Harrison 1999; Arnold et al. 1998), yet it is consistently associated with dysfunction of cortical circuits, particularly in the prefrontal cortex (PFC), with impairments in glutamatergic neurons (Glantz and Lewis 2000; Glausier and Lewis 2013) and γ-aminobutyric acid (GABA)-mediated inhibition (Lewis et al. 2005). Abnormalities in GABAergic interneurons are among the most reproducible neurobiological findings in SCZ, with the somatostatin (Sst) and parvalbumin (Pvalb) interneuron subclasses most frequently implicated (Dienel et al. 2023; Batiuk et al. 2022; Hashimoto et al. 2008; Mulvey et al. 2025; Arbabi et al. 2025).

Whether these interneurons are also fewer in number, however, has remained contested. *In situ* studies report lower SST and GABA-synthetic transcripts per neuron without a detectable change in neuron density (Dienel et al. 2023; Schahram Akbarian 1995; Hashimoto et al. 2008), whereas histological counts and bulk-tissue deconvolution report or infer fewer interneurons (Toker et al. 2018; Kiss et al. 2026; Batiuk et al. 2022; Benes 1991; Beasley et al. 2002). Part of the difficulty in reconciling these alternatives is methodological, because in marker-based cell counting, a surviving neuron expressing undetectable levels of its identifying marker is indistinguishable from one that is truly absent (Dienel et al. 2023). The evidence therefore leaves two possibilities, (1) that interneuron number is preserved and only their molecular state is altered, or (2) that some interneurons are altered and also fewer in number. Resolving this question requires the ability to distinguish closely related inhibitory cell types, because a change confined to a specific subset of neurons may be masked or diluted when cells are analyzed only at a coarse level.

Recent single-nucleus transcriptomic atlases now provide taxonomies of human neocortex at unprecedented granularity. The Seattle Alzheimer’s Disease Brain Cell Atlas (SEA-AD) expands the conventional ∼24 cortical subclasses (e.g., Sst) into 137 finer “supertypes” (e.g., Sst_25), many of which are spatially restricted to particular cortical layers (Gabitto et al. 2024). Corresponding Patch-seq datasets also enable linking these transcriptomic types to their morphoelectric and laminar properties (B. R. Lee et al. 2023), such that a transcriptomic label (i.e. any given supertype) can be tied to multi-modal cellular phenotypes. In parallel, the rapid growth of publicly available post-mortem SCZ single-nucleus RNA-sequencing (snRNA-seq) datasets creates an opportunity to test cell type-specific hypotheses at scale (Ruzicka et al. 2024; Ling et al. 2024), thus addressing the problems associated with small sample sizes and dataset heterogeneity that have limited individual studies.

Here we present the largest case-control single-cell analysis of post-mortem brain in SCZ to date, encompassing eight datasets and almost 500 donors, to address three questions about cortical inhibitory interneuron pathology. First, do interneurons persist in an altered molecular state, are some of them lost, or both? Second, if their numbers change, which specific cell types are most affected? Third, what distinguishes the vulnerable cell types — concentrated SCZ genetic risk, and/or defined morphoelectric properties? To resolve these questions at a granularity that separates closely related cell types, and with statistical power that individual post-mortem studies have lacked, we integrate cell type-specific differential expression and differential abundance across seven snRNA-seq datasets, replicate both the expression and abundance findings in an independent Xenium spatial transcriptomics dataset, and connect the implicated supertypes to SCZ common-variant genetic risk and to their physiological and morphological characteristics.

## Results

### A unified cell-type taxonomy organizes cells across seven snRNA-seq datasets and spatial transcriptomics sections

To examine how cortical cell types change in SCZ, we compiled seven post-mortem snRNA-seq datasets with tissue collected from the prefrontal cortex, totaling 469 donors (Table 1; 298 neurotypical controls and 171 SCZ cases; approximately 2.4 million nuclei). We mapped every nucleus onto the SEA-AD taxonomy of neocortical cell types, defined from neurotypical donors (Gabitto et al. 2024), assigning each nucleus to one of 24 subclasses and, within each subclass, to one of 137 finer supertypes (Fig. 1a; Methods). Throughout, we follow the convention of SEA-AD taxonomic nomenclature: class (e.g., GABAergic), subclass (e.g., Sst cells), and supertype (e.g., Sst_25 cells) denote successively finer levels of the same taxonomy, and “interneuron” refers to GABAergic neurons. Label transfer was run separately within each dataset against the SEA-AD reference (Methods), so cell type labels are directly comparable across datasets and provide the common framework for the meta-analyses that follow. In a joint embedding, nuclei grouped by cell type rather than by dataset of origin or diagnosis (Fig. 1b–d) and cell types expressed their expected marker genes (Fig. S1).

**Fig. 1.**
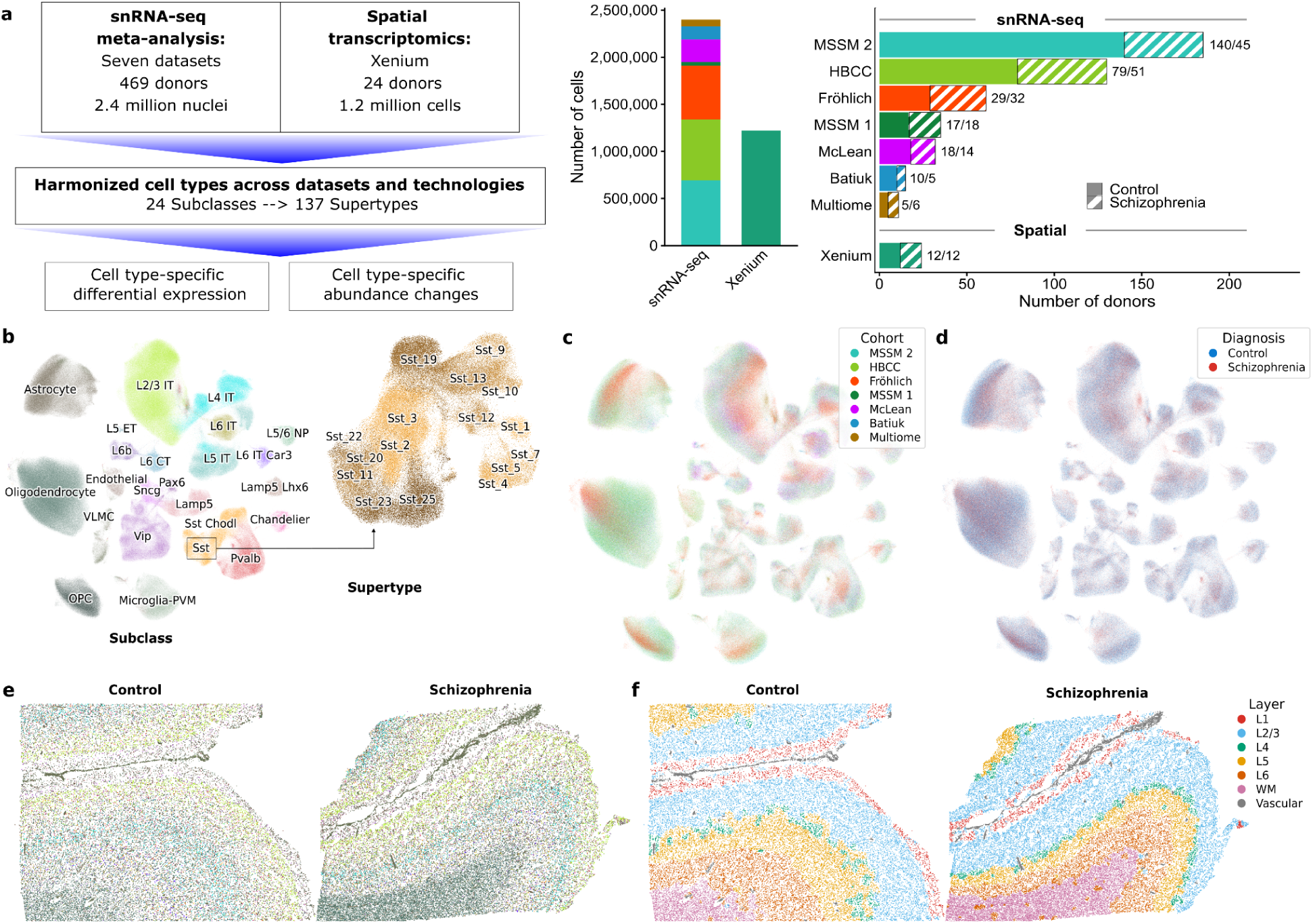
Study design, cross-cohort snRNA-seq harmonization, and Xenium spatial annotation. (**a**) Left, flowchart illustrating seven-dataset snRNA-seq meta-analysis and spatial transcriptomics analysis workflow. Middle, number of nuclei (or cells) profiled per dataset and platform. Right, donors sampled per dataset, split into neurotypical controls (solid) and SCZ cases (hatched); bar plot annotations denote control/SCZ donor counts. (**b**) UMAP of the integrated snRNA-seq datasets colored by cell subclass (left); nuclei of the Sst subclass colored by their 16 constituent supertypes (right). (**c, d**) The same integrated UMAP embedding as in b colored by (**c**) dataset of origin and (**d**) diagnosis. (**e, f**) Representative Xenium sections, with each cell colored by annotated subclass (**e**) and inferred cortical layer (**f**).

**Table 1.** Datasets and donor demographics. Post-mortem donor counts, age, and brain region for every SCZ case/control dataset analyzed in this study. The upper block lists the seven snRNA-seq datasets meta-analyzed for differential expression and abundance, followed by their pooled totals (Meta-analysis row; 298 control and 171 SCZ donors, 469 across the seven datasets). The lower block lists the DLPFC Xenium spatial transcriptomics dataset used for orthogonal replication. Only donors younger than 70 years at death were included (Methods). Additional per donor metadata is available in Supplemental Table T1. Abbreviations: CON, control; SCZ, schizophrenia; DLPFC, dorsolateral prefrontal cortex; OFC, orbitofrontal cortex; snRNA-seq, single-nucleus RNA sequencing.

| Dataset | Data type | CON (N) | SCZ (N) | Mean age (SD) | Male (%) | Region | Brain bank | Citation |
| --- | --- | --- | --- | --- | --- | --- | --- | --- |
| Batiuk | snRNA-seq | 10 | 5 | 58.7 (5.3) | 60 | DLPFC | Netherlands Brain Bank; Oxford Brain Bank; Human Brain Tissue Bank, Semmelweis University; Newcastle Brain Bank | Batiuk et al. 2022 |
| McLean | snRNA-seq | 18 | 14 | 53.4 (10.6) | 53.1 | PFC | Harvard Brain Tissue Resource Center, McLean Hospital | Ruzicka et al. 2024 |
| MSSM 1 | snRNA-seq | 17 | 18 | 55.8 (12.2) | 77.1 | PFC | Mount Sinai NIH NeuroBioBank | Ruzicka et al. 2024 |
| Multiome | snRNA-seq | 5 | 6 | 43.1 (13.1) | 63.6 | DLPFC | NIMH Human Brain Collection Core | Emani et al. 2024 |
| Fröhlich | snRNA-seq | 29 | 32 | 52.9 (9.8) | 67.2 | OFC | New South Wales Brain Tissue Resource Centre, University of Sydney | Fröhlich et al. 2024 |
| HBCC | snRNA-seq | 79 | 51 | 43.7 (12.7) | 60 | DLPFC | NIMH Human Brain Collection Core | D. Lee et al. 2024 |
| MSSM 2 | snRNA-seq | 140 | 45 | 56.2 (11.1) | 71.4 | DLPFC | Mount Sinai NIH NeuroBioBank | D. Lee et al. 2024 |
| Meta-analysis | snRNA-seq | 298 | 171 | 51.8 (12.6) | 66.3 | PFC | – | – |
| Xenium (LIBD) | Xenium | 12 | 12 | 47.9 (8.9) | 45.8 | DLPFC | Lieber Institute for Brain Development | Kwon et al. 2026 |

To test whether the snRNA-seq findings replicate in a spatially resolved data modality, we added an independent Xenium spatial transcriptomics dataset of 24 dorsolateral prefrontal cortex (DLPFC) sections (12 controls and 12 SCZ cases), profiled with a custom 300-gene panel of cell type markers and genes previously reported as differentially expressed in SCZ (Kwon et al. 2026). We assigned each Xenium cell to the same taxonomy used for the snRNA-seq data, using custom approaches for cell typing and for inferring cortical depth and layer (Supplementary Methods, Figs S2-3), after which 1.2 million cells remained following quality control (Fig. 1e,f).

## Somatostatin interneurons show a reduction of *SST* mRNA that is consistent across datasets and platforms

We first asked whether the canonical inhibitory markers most often implicated in SCZ, *SST* and *PVALB*, are expressed at lower transcript levels within the interneurons that they define. We therefore compared *SST* mRNA in Sst interneurons and *PVALB* mRNA in Pvalb interneurons, between SCZ cases and controls in each of the seven snRNA-seq datasets (adjusting for age, sex and post-mortem interval; Methods). We then pooled the seven estimates by meta-analysis and sought independent replication in Xenium.

Within Sst interneurons, *SST* mRNA was significantly reduced in SCZ (Fig. 2a,b, meta-analyzed log_2_ fold change [FC] = −0.46, ∼27% lower SST mRNA in SCZ, *P* = 6.3 × 10^-4^, False Discovery Rate [FDR] = 0.048 transcriptome-wide). Because expression was compared within Sst cells, this reduction reflects lower *SST* mRNA per Sst cell rather than fewer Sst cells. This comparison does not depend on detecting *SST* in each cell as Sst interneurons were identified from transcriptome-wide profiles (Methods), so nuclei with no detectable *SST* transcript, which were more frequent in SCZ (41% versus 33%, *P* = 2 × 10^-4^), were nonetheless classified as Sst and retained. The fold change estimate was negative in six of the seven datasets and nominally significant in three (i.e., unadjusted *P* < 0.05), with low heterogeneity (I² = 16.6%) in the meta-analysis. Moreover, the Xenium spatial transcriptomics dataset showed a similar overall reduction (Fig. 2b–d, log_2_FC = −0.33, *P* = 0.052). The *SST* reduction was also present within individual Sst supertypes (Fig S4; negative log_2_FC in 11 of the 13 tested supertypes), suggesting a subclass-wide rather than supertype-specific deficit. *SST* was one of 343 genes differentially expressed (DE) in Sst cells at FDR < 0.10 (Fig. 2a; 196 down, 147 up; Supplemental Table T2), the strongest of which were *NAT16* (down-regulated) and *SMAD1* (up-regulated).

**Fig. 2.**
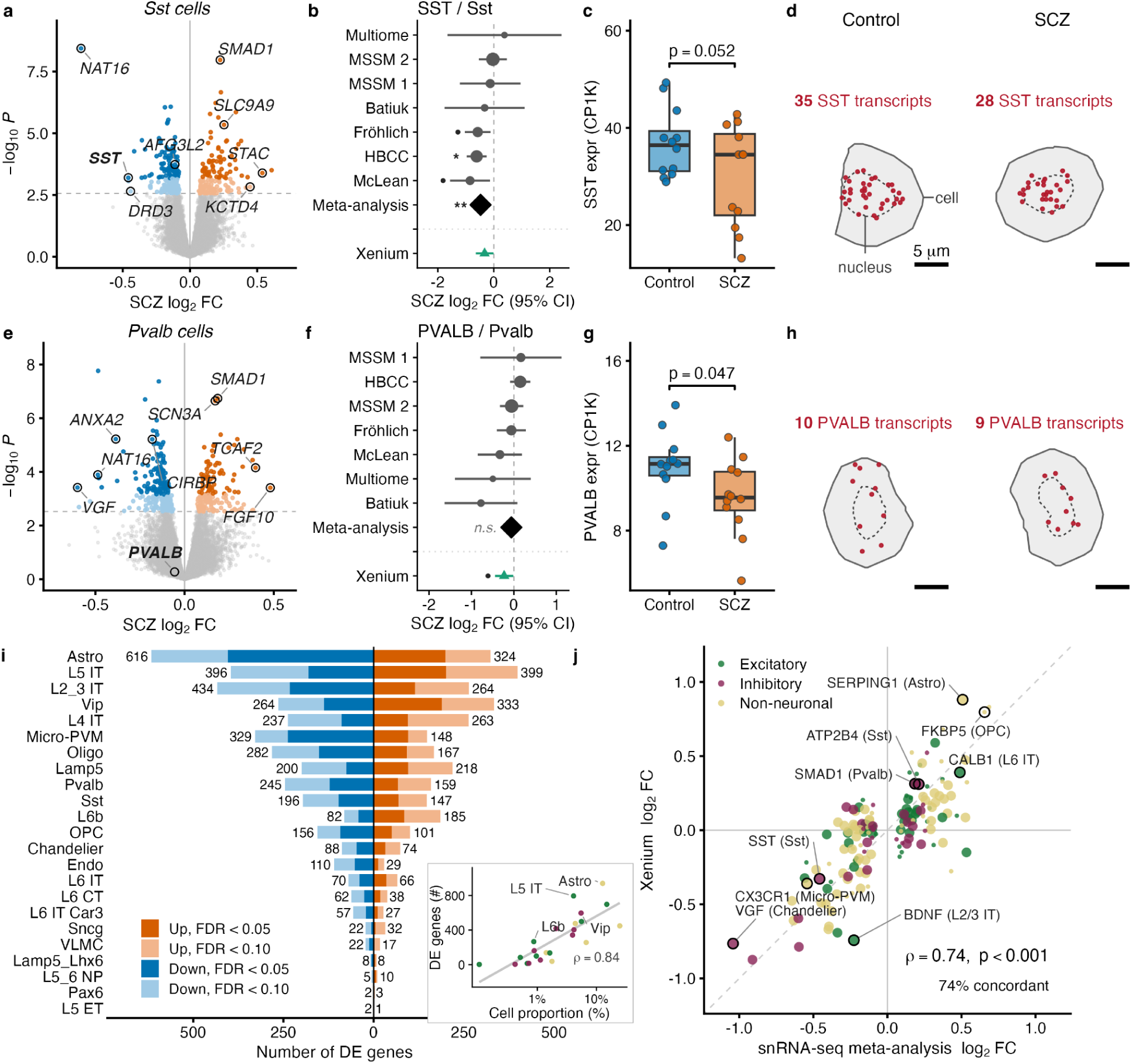
Cell-type–specific differential expression in schizophrenia. (**a–d**) Differential expression (DE) in Sst interneurons. (**a**) Volcano plot of SCZ log_2_ fold change for all genes tested in Sst cells in the seven-dataset snRNA-seq meta-analysis. Selected genes labeled, *SST* in bold. (**b**) Forest plot of *SST* mRNA dysregulation in each of seven snRNA-seq datasets (gray circles; whiskers, 95% CI; dot size proportional to donor count), pooled by random-effects meta-analysis (black diamond), and in Xenium (green triangle). (**c**) *SST* mRNA expression in Xenium Sst cells per donor (counts per 1,000 transcripts, CP1K), 12 control versus 12 SCZ donors. (**d**) One representative Xenium Sst cell per group (cells chosen closest to each group’s median *SST* transcript density); solid and dashed lines indicate cell and nucleus boundaries respectively and colored dots indicate *SST* transcripts; scale bar, 5 µm. (**e–h**) Differential expression in Pvalb interneurons, as in a-d. (**i**) Number of DE genes per subclass in the meta-analysis, shaded by FDR tier; DE counts at FDR < 0.10 indicated at bar ends. Inset, DE genes per subclass versus the subclass’s mean proportion of nuclei per donor (log axis); Spearman ρ. (**j**) Xenium versus meta-analysis log_2_ fold change for the 167 gene × cell type pairs DE in the meta-analysis (FDR < 0.10) and measured on the Xenium panel; larger points, FDR < 0.05; dashed line, identity; selected subclass-gene pairs labeled. Points in the inset of i and in j are colored by cell class (green, excitatory; purple, inhibitory; yellow, non-neuronal). Significance throughout: ***FDR < 0.01, **FDR < 0.05, *FDR < 0.10; •, nominal *P* < 0.05; n.s., not significant.

The second canonical marker, *PVALB*, was not DE within Pvalb interneurons in the snRNA-seq meta-analysis (Fig. 2e,f, log_2_FC = −0.06, *P* = 0.54, FDR = 0.85), nor in any individual snRNA-seq dataset. In the Xenium dataset, *PVALB* was nominally reduced in Pvalb interneurons (Fig. 2f–h; log_2_FC = −0.23, *P* = 0.047), a discrepancy consistent with a partly cytoplasmic *PVALB* signal that Xenium captures but nucleus-restricted snRNA-seq may not (see Discussion). Beyond *PVALB* itself, 404 genes were DE in Pvalb cells at FDR < 0.10 (Fig. 2e; 245 down, 159 up), the strongest of which were *SMAD1* and *SCN3A* (both up-regulated).

Expanding this analysis transcriptome-wide to all cell types at the subclass resolution, the meta-analysis identified 6,898 gene x cell type pairs that were DE at FDR < 0.10, with more genes down- than up-regulated (Fig. 2i; 3,885 down, 3,013 up). The number of DE genes observed in each subclass strongly tracked the overall abundance of the subclass in the snRNA-seq datasets (Fig. 2i inset; Spearman ρ = 0.84), suggesting greater power in more abundant cell types. For example, astrocytes, among the most abundant of subclasses, had the most DE genes (940), whereas rarer subclasses, such as L5 ET excitatory and Pax6 GABAergic cells, had five or fewer. Because abundant subclasses contribute more nuclei per donor and therefore likely have greater statistical power to identify DE, we do not read the raw DE-gene count as a measure of how strongly a cell type is affected in SCZ. The same dependence on nuclei number left supertype-level DE underpowered for most supertypes (Fig. S4, full results in Supplemental Table T3).

We next asked whether the cell type-specific DE observed in snRNA-seq reproduced in Xenium. Of the gene x subclass DE pairs in the meta-analysis (FDR < 0.10), 167 overlapped genes measured on the 300-gene Xenium panel. Across these pairs, the Xenium and snRNA-seq meta-analysis fold changes were positively correlated and agreed in direction for 74% (Fig. 2j; Spearman ρ = 0.74). For example, *FKBP5* was up-regulated in OPCs on both platforms (meta-analysis log_2_FC = +0.66, Xenium log_2_FC = +0.80) and *SERPING1* in astrocytes, whereas *VGF* in Chandelier cells, *BDNF* in L2/3 IT cells, and *CX3CR1* in Microglia-PVM cells were consistently down-regulated.

### Upper-layer Sst supertypes are less abundant and L6b excitatory supertypes more abundant in schizophrenia

Next, we asked whether cell type abundances also change in SCZ; for example, whether the disorder is accompanied by fewer Sst or Pvalb interneurons. We compared the donor-level proportion of each cell type between SCZ cases and controls in each of the seven snRNA-seq datasets (using crumblr (Hoffman and Roussos 2025) adjusting for age, sex and post-mortem interval; Methods). We then pooled the dataset independent estimates by fixed-effect meta-analysis and sought independent replication in Xenium.

Abundance changes were confined to only a few neuronal supertypes (Fig. 3a; Supplemental Table T4). Five Sst supertypes were less abundant in SCZ (hereafter, depleted); four at FDR < 0.05 (Sst_2, Sst_22, Sst_25 and Sst_3; β = −0.19 to −0.27, *P* ≤ 2.6 × 10^-3^, FDR 0.0009 to 0.046) and Sst_20 at FDR < 0.20 (β = −0.17, *P* = 0.019, FDR = 0.19). Two layer 6b excitatory supertypes were more abundant in SCZ (L6b_1, β = +0.48, *P* = 1.9 × 10^-6^, FDR = 2 × 10^-4^; L6b_4, β = +0.26, *P* = 5.9 × 10^-4^, FDR = 0.022). No Pvalb or non-neuronal supertype changed in abundance in SCZ at FDR < 0.10 (Fig. S5). Trend-level increases, at FDR < 0.20, were seen for Pvalb_14 and three excitatory supertypes (L2/3 IT_7, L6 CT_1, L5/6 NP_4).

**Fig. 3.**
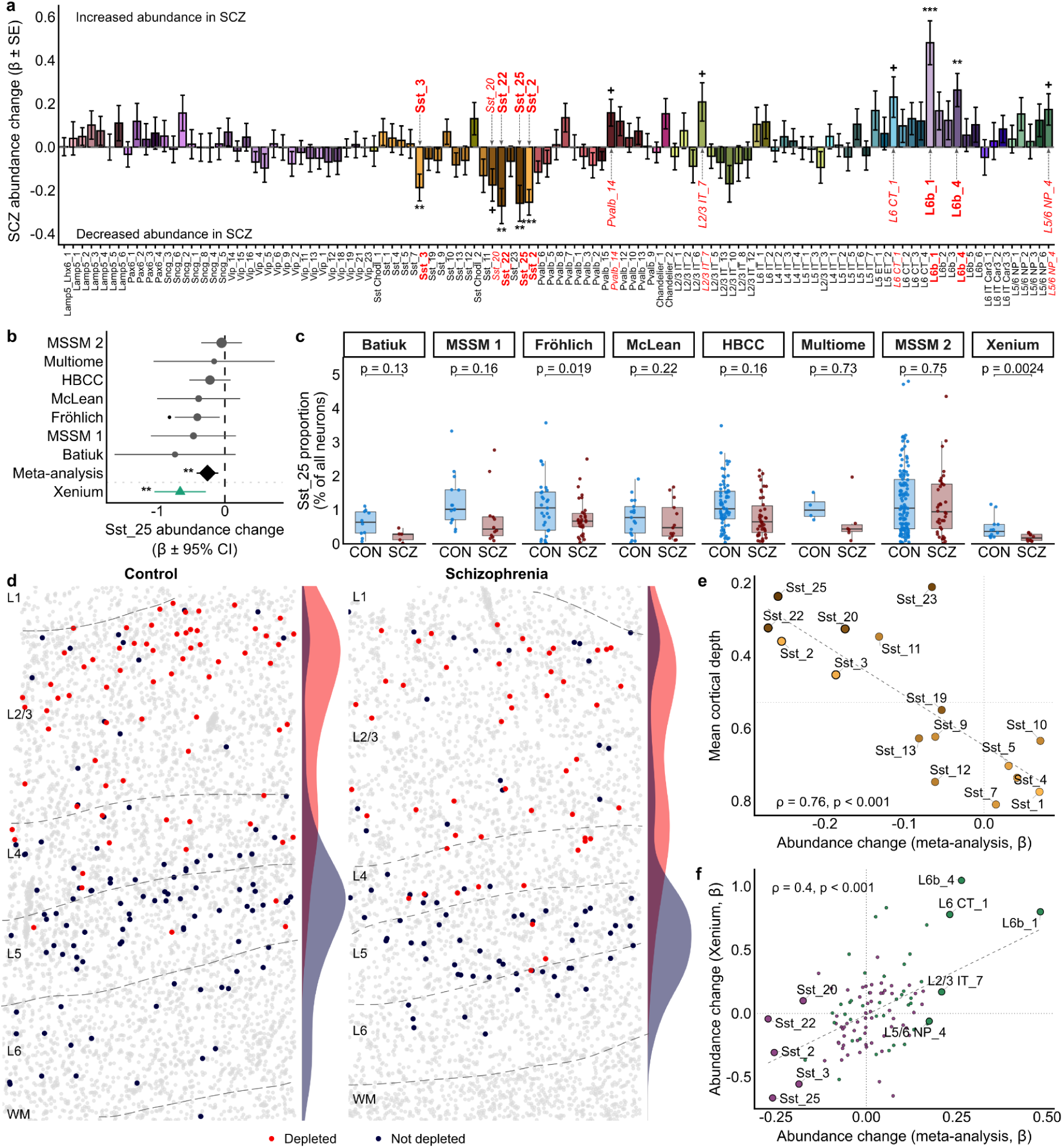
Differential abundance of neuronal cell types in schizophrenia. (**a**) Effect of SCZ on the relative abundance of each of 109 neuronal supertypes, from the seven-dataset snRNA-seq meta-analysis (crumblr β ± SE). Negative bars denote lower abundance (depletion) in SCZ. Bars colored by supertype; supertype names in bold red, FDR < 0.10; in italic red, FDR < 0.20. (**b**) Forest plot of the Sst_25 SCZ abundance change in each of the seven snRNA-seq datasets (gray circles; whiskers, 95% CI; circle size proportional to donor count), pooled by fixed-effects meta-analysis (black diamond), and in Xenium (green triangle). (**c**) Sst_25 as a percentage of all neurons per donor in each snRNA-seq dataset and in Xenium, control (blue) versus SCZ (red); unadjusted P values. (**d**) Exemplar control and SCZ Xenium sections; Sst cells colored by whether their supertype is depleted in a (red; Sst_2, Sst_3, Sst_20, Sst_22, Sst_25) or not (navy), other cells gray; curves at right, depth distribution of each Sst group. (**e**) Mean cortical depth from Xenium (0 = pia, 1 = white matter; pia at top) versus snRNA-seq meta-analysis SCZ abundance change for the 16 Sst supertypes. (**f**) Xenium versus snRNA-seq abundance changes for matched neuronal supertypes (purple, inhibitory; green, excitatory); supertypes highlighted in red in a are enlarged and labeled; dashed line denotes identity. Significance throughout: ***FDR < 0.01, **FDR < 0.05, *FDR < 0.10, +FDR 0.10–0.20, • nominal *P* < 0.05.

Individual datasets were generally underpowered to detect compositional changes, although effects were consistent in direction across datasets (Fig. S6). Sst_25, for example, had a per-donor proportion only modestly lower in SCZ, reaching nominal significance in one dataset (Fig. 3b,c; Fröhlich, *P* = 0.019). Pooling across the seven datasets gave an estimated 27% reduction (95% CI: 14-39%), as Sst_25 made up 1.01% of neurons in controls versus 0.74% in SCZ cases. These overall abundance effects were robust to the pooling strategy (fixed-effect or random-effects meta-analysis, or analyzing all donors together jointly in a single model) and to leaving out any single dataset (Fig. S6). Because the expression changes observed in Fig. 2 could in principle bias cell typing (Dienel et al. 2023), we re-annotated every nucleus after excluding all 343 genes DE in Sst interneurons at FDR < 0.10, including *SST* mRNA itself; this re-annotation recovered the same abundance changes (Fig. S7; all five Sst supertypes depleted at FDR < 0.12), so the observed depletion is unlikely to be an artifact of expression-driven mislabeling (see Discussion).

We next asked whether these abundance changes replicate in cells observed *in situ*, measured using Xenium spatial transcriptomics, applying the same analysis to the 24 Xenium DLPFC sections. Three of the five depleted Sst supertypes were also significantly depleted in these tissue sections (Fig. 3b,c,f; Sst_25, Sst_3 and Sst_2, β = −0.31 to −0.66, all *P* < 0.03), and both L6b supertypes were more abundant (β = +0.80 and +1.05, both *P* < 0.01). Sst_22 and Sst_20 were not replicated, which may reflect the limited ability of the 300-gene panel to resolve these supertypes (Fig. S2; Supplementary Methods). In the Xenium sections, cells of the depleted Sst supertypes were visibly concentrated in superficial cortical layers (Fig. 3d). Relating each supertype’s mean cortical depth in Xenium to its abundance change in the snRNA-seq meta-analysis, we found that more superficial Sst supertypes were more depleted in SCZ (Fig. 3e; Spearman ρ = 0.76, *P* < 0.001). Across the 106 neuronal supertypes present on both platforms, per-supertype abundance changes were positively, if modestly, correlated (Fig. 3f; Spearman ρ = 0.40, *P* < 0.001).

We then asked whether the transcriptional state of Sst interneurons differs between the supertypes that are depleted and those that persist. We grouped the 16 Sst supertypes by their depletion status in Fig. 3a into three broad groups: depleted (FDR < 0.20; n = 5), intermediate (reduced abundance, but non-significant; n = 6) and non-depleted (n = 5). We repeated the differential expression analysis of Fig. 2 followed by gene set enrichment analysis within each group (Fig. S8, Methods). We found that *SST* and *VGF* mRNA and synaptic gene sets were generally reduced in all three groups. In contrast, gene sets for cytosolic translation were downregulated only in the depleted and intermediate groups, and gene sets for oxidative phosphorylation were downregulated in the depleted group alone. Although changes at the level of individual genes were modest, these results suggest that while aspects of transcriptional dysregulation are broadly shared across Sst interneurons, suppression of protein synthesis and oxidative phosphorylation is largely confined to the supertypes undergoing depletion.

### Schizophrenia genetic risk converges on *HCN1*- and *CALB1*-expressing upper-layer Sst supertypes that are also depleted in Alzheimer’s disease

Finally, we asked whether the depletion of upper-layer Sst supertypes is etiologically relevant to SCZ, or instead a downstream consequence of the disorder or its treatment. Because common-variant risk is germline and precedes the disorder, the cell types it implicates are more plausibly part of the causal pathway than a consequence (Bryois et al. 2020; Finucane et al. 2018; Skene et al. 2018). Recent work has shown that cortical Sst subtypes carry the strongest SCZ genetic association of any brain cell type in neurotypical atlases (Duncan et al. 2025); however, these approaches cannot show whether the same subtypes are also most altered in disease. We therefore asked whether the Sst supertypes depleted in our case-control data are also those in which SCZ risk genes (Bigdeli et al. 2026) are most specifically expressed. We applied the enrichment approach of Duncan et al. (2025) to the same SEA-AD supertypes, taking each supertype’s expression profile from the region-matched neurotypical DLPFC reference (Methods; Fig. S9).

Across the 16 Sst supertypes, those whose specific genes carried the most SCZ common-variant association, including Sst_2, Sst_3 and Sst_20, tended also to be the most depleted (Fig. 4a; Spearman ρ = 0.55, *P* = 0.031; robust to GWAS and expression reference, Fig. S10). For Sst_2, the most enriched and among the most depleted supertypes, we asked which Sst_2-specific genes carry the SCZ association (Fig. 4b; Methods). Among them was *HCN1* (top 10% of Sst_2 specificity; MAGMA −log_10_ P = 10.9), the only protein-coding gene within 300 kb of a genome-wide significant SCZ locus whose fine-mapped credible set sits at the gene’s 3′ end (Fig. 4c), and previously prioritized as the likely causal gene there by our PsyOPS method (Wainberg et al. 2022). *HCN1* encodes a subunit of the channels carrying the hyperpolarization-activated current I_h, which dampens excitability and produces a voltage sag at hyperpolarized potentials. In Patch-seq recordings of 129 human neocortical Sst interneurons (B. R. Lee et al. 2023), sag was larger in supertypes with higher *HCN1* expression (Fig. 4d; ρ = 0.65, *P* = 0.008; example traces in Fig. 4f). These recordings come from neurosurgical tissue of individuals without SCZ, describing the intrinsic physiology of the vulnerable supertypes and not a change in SCZ, which remains untested.

**Fig. 4.**
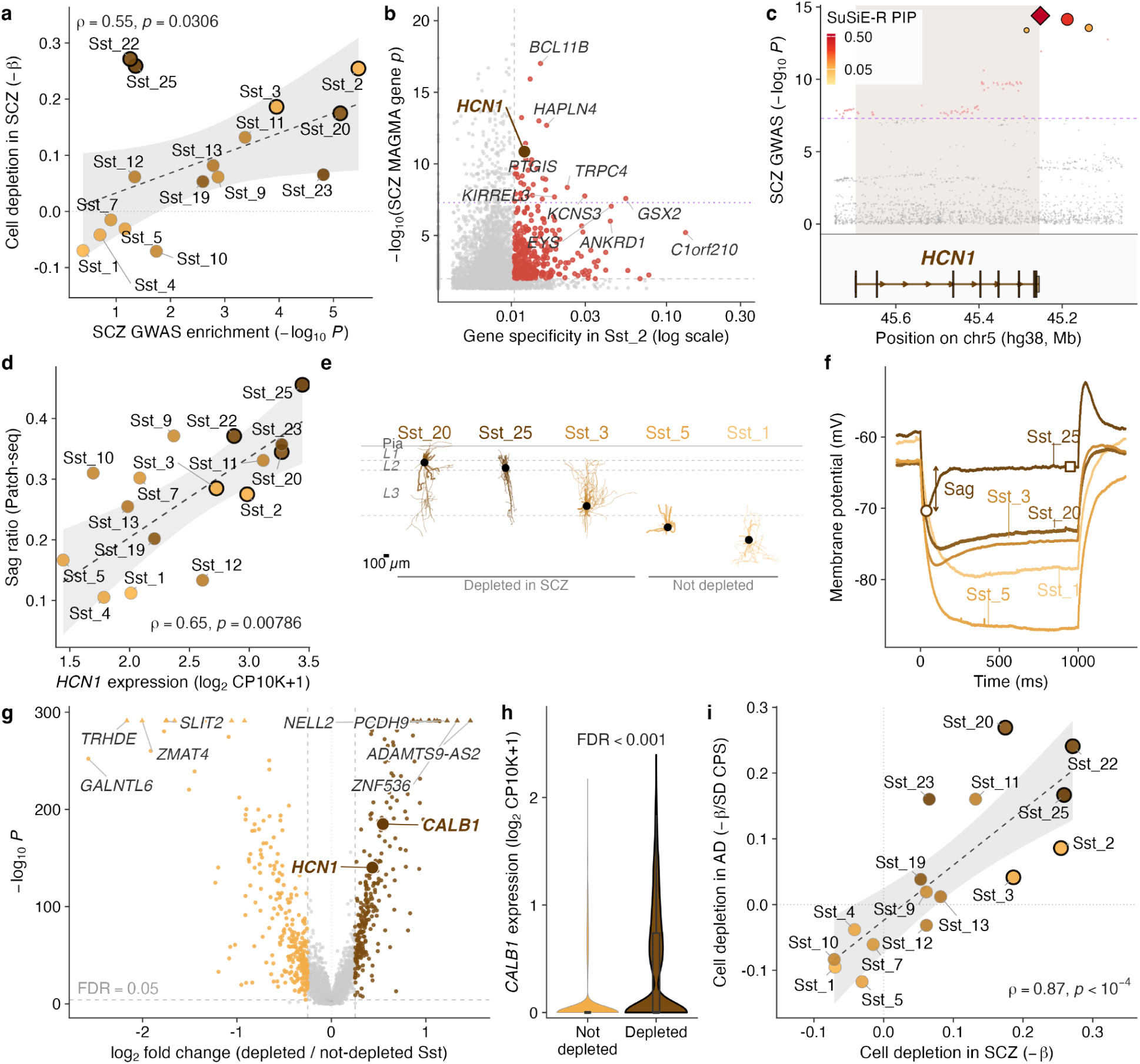
Genetic risk, HCN1-dependent physiology, and cross-disorder vulnerability of depleted upper-layer Sst supertypes. (**a**) SCZ GWAS enrichment per supertype (−log_10_ P) versus compositional depletion in SCZ (−β from Fig. 3a). In a, and throughout the figure, each point is one of the 16 Sst supertypes; thick outlines mark the five depleted in SCZ; dashed lines, linear fits with 95% confidence bands; ρ, Spearman. All neurotypical expression data are from the SEA-AD DLPFC snRNA-seq reference, which shares the supertype taxonomy used throughout. (**b**) Expression specificity for Sst_2 versus gene-level SCZ genetic association (−log_10_ P). Vertical dashed line, top 10% most Sst_2-specific genes; horizontal lines, gene-level FDR 0.05 (gray) and genome-wide significance (purple); genes passing both cut-offs in red; *HCN1* highlighted. (**c**) SCZ GWAS association across the HCN1 locus; fine-mapped variants enlarged and colored by posterior inclusion probability (PIP); the variant set sits at the 3′ end of *HCN1*, with the lead variant rs10035564, 2.5 kb beyond the transcript end. (**d**) Mean *HCN1* expression per supertype versus mean voltage-sag ratio from supertype-matched human Patch-seq data. (**e**) Exemplar Patch-seq morphological reconstructions per supertype, ordered by soma depth (black dot), for depleted (Sst_20, Sst_25, Sst_3) and not-depleted supertypes in SCZ (Sst_5, Sst_1); dashed lines, layer boundaries; scale bar, 100 µm. (**f**) Voltage responses of the cells in e to a hyperpolarizing current step (0–1000 ms); sag marked between voltage minimum (circle) and steady state (square). (**g**) Differential expression between depleted and not-depleted Sst supertypes in the SEA-AD neurotypical DLPFC reference (3 donors); brown, higher in depleted; light orange, higher in not-depleted; dashed lines, |log_2_FC| = 0.25 and FDR = 0.05; triangles, P values beyond the y-axis range; *HCN1* and *CALB1* labeled. (**h**) Per-cell *CALB1* expression by depletion group; cell-level Wilcoxon test, FDR < 0.001. (**i**) Depletion in SCZ (−β, as in a) versus depletion in AD, as indexed by decline along the SEA-AD AD pseudo-progression score (CPS) in DLPFC (−β per SD of CPS; 80 donors).

We next asked what other genes distinguish the depleted supertypes, and identified 580 genes differentially expressed between depleted and not-depleted Sst supertypes in the SEA-AD neurotypical reference (FDR < 0.05, |log_2_FC| > 0.25; Fig. 4g). *HCN1* was among them, more highly expressed in the depleted supertypes (log_2_FC = +0.43). As was *CALB1*, encoding the calcium-binding protein calbindin (Fig. 4h; log_2_FC = +0.55, FDR < 0.001). Calbindin marks several upper-layer Sst supertypes, including double-bouquet cells, a morphological type with no clear rodent counterpart (DeFelipe et al. 2006); among the depleted supertypes, Sst_25 in particular largely corresponds to double-bouquet cells (Gabitto et al. 2024; Zaitsev et al. 2009). Reconstructed exemplars of the depleted supertypes sit in layers 2/3 with narrow, vertically oriented axonal bundles and show pronounced sag, whereas exemplars of not-depleted supertypes sit deeper, are multipolar, and show little sag (Fig. 4e,f).

We also asked whether the Sst supertypes depleted in SCZ are depleted in Alzheimer’s disease, where Sst subtypes have likewise been reported to be lost (Gabitto et al. 2024; Mathys et al. 2024; Travaglini et al. 2026; Consens et al. 2022). The SEA-AD resource (Gabitto et al. 2024) profiled DLPFC and MTG by snRNA-seq from a shared set of 80 aged donors spanning the AD neuropathological continuum, and assigned each donor a continuous pseudo-progression score (CPS). SEA-AD uses the same supertype taxonomy as this study, so we could directly relate each supertype’s SCZ differential abundance (Fig. 3a) to its change along the AD pseudo-progression score. In DLPFC, the SCZ and AD effects agreed in direction for 15 of 16 Sst supertypes, and the SCZ-depleted supertypes were among those declining most as AD advances (Fig. 4i; Spearman ρ = 0.87, P < 10^-4^, n = 16; MTG comparison in Fig. S10). The same upper-layer Sst supertypes are therefore depleted in two disorders with distinct etiologies and neuropathologies.

## Discussion

Whether cortical inhibitory interneurons in SCZ are fewer in number, in addition to being molecularly altered, has been debated for over two decades. The dominant view, built on *in situ* measurements of transcripts per neuron and of neuron density, is that interneurons are altered in their molecular state without detectable cell loss (Volk et al. 2000; Hashimoto et al. 2003; Dienel et al. 2023, 2022). In contrast, a smaller body of histological and bulk-tissue deconvolution work reports fewer interneurons (Benes 1991; Beasley et al. 2002; Toker et al. 2018; Batiuk et al. 2022; Kiss et al. 2026). These accounts were never mutually exclusive, and our data, a seven-dataset snRNA-seq meta-analysis with spatial replication, reconcile them. The reduction of *SST* and other transcripts within Sst neurons is subclass-wide, and it co-occurs with the depletion (reduced abundance) of specific upper-layer Sst supertypes. Beyond establishing the depletion itself, we show that the depleted supertypes show concentrated SCZ genetic risk, are marked by high *HCN1* expression and HCN-dependent physiology, downregulate translation and energy-metabolism programs in SCZ, and are also among the earliest neurons lost in Alzheimer’s disease.

Because SCZ alters the very transcripts that define cell identity, an apparently depleted cell type could be truly absent or merely unrecognizable (identity erosion), the confound that Dienel et al. (2023) described for marker-based counting, here across the whole transcriptome. Three observations argue against erosion. First, mirroring the solution of Dienel et al. (2023) to identify neurons through markers the disorder leaves intact, we repeated cell typing after excluding all genes detectably altered in Sst interneurons, including *SST* itself, and recovered the same depletion. Mislabeling would therefore require expression shifts each too small to detect, yet collectively severe enough to erase supertype identity. Second, label transfer assigns every cell to its best-matching type, so eroded cells would be relabeled rather than uncounted, yet no other interneuron supertype increased in abundance at FDR < 0.10. Third, we found similar Sst supertype depletion *in situ* in a different donor cohort, using a separate technology (Xenium), gene panel, and cell classification approach. We therefore regard a genuine reduction in cell number in SCZ as the most parsimonious interpretation. Either account entails substantial disruption of the same neurons, so the remaining ambiguity concerns the mechanism of depletion rather than which cells are affected.

Beyond the broad reduction of *SST* mRNA, we observed that the altered molecular state of Sst interneurons had two major components. One component was broadly shared across Sst supertypes regardless of depletion status, including reductions in *VGF* and synaptic gene sets, and echoing the Sst transcriptional pathology reported across psychiatric disorders (Newton et al. 2022). By contrast, suppression of protein-synthesis and oxidative-phosphorylation programs was graded by depletion status and was strongest in the supertypes reduced in number, consistent with rodent work in which chronic stress suppresses translation selectively in Sst neurons (Lin and Sibille 2015; Tomoda et al. 2022).

Laminar-specific interneuron pathology in SCZ was described well before transcriptomic taxonomies existed. Reduced density of calbindin-immunoreactive interneurons, cells concentrated in superficial cortical layers and especially in layer 2, was reported in SCZ prefrontal cortex (Beasley et al. 2002) and planum temporale (Chance et al. 2005) more than twenty years ago; however, the identity of the affected cells could only be inferred from calbindin immunoreactivity, not demonstrated. Our findings plausibly attribute these prior observations to named Sst supertypes, as our depleted cells concentrate in upper cortical layers and the depleted supertypes preferentially express the gene encoding calbindin, *CALB1*.

Across Sst supertypes, genetic risk enrichment tracked the degree of depletion. Because germline risk precedes both disease onset and medication exposure, this convergence favors depletion being part of the disorder’s primary pathology rather than a consequence of the disorder or its treatment (Bryois et al. 2020; Finucane et al. 2018; Skene et al. 2018). Among the genes carrying this signal, *HCN1* especially stood out as it is the likely causal gene at its SCZ risk locus (Wainberg et al. 2022) and is more highly expressed in depleted supertypes. Consistent with HCN1 function, the depleted supertypes display pronounced HCN-dependent voltage sag in Patch-seq recordings from neurotypical human cortex, a property that is itself markedly elevated in human relative to mouse interneurons (B. R. Lee et al. 2023). High expression of SCZ risk genes such as *HCN1* thus appears to be part of these cells’ baseline identity, accompanying a distinctive physiology that shapes how they integrate and time synaptic input. Disruption of that processing is therefore a plausible route to the deficits in information processing and cognition that are among the most disabling features of SCZ (Kahn et al. 2015).

Intriguingly, the Sst supertypes we find depleted in SCZ are also among the earliest neuronal populations lost as Alzheimer’s disease advances (Gabitto et al. 2024. Convergence of two disorders with distinct etiologies and largely unshared genetic risk factors on the same small set of neurons suggests a vulnerability intrinsic to the cells themselves (Saxena and Caroni 2011). We hypothesize that the specializations making these cells distinctive also render them especially fragile. For example, in addition to their high-sag physiology, the depleted supertypes include Sst_25, which largely corresponds to the double-bouquet cell (Gabitto et al. 2024), a morphology first described by Ramón y Cajal in human cortex (Ramón y Cajal 1899; DeFelipe et al. 2006) and lacking any clear rodent counterpart (Ballesteros-Yáñez et al. 2006; Raghanti 2010). The depleted supertypes thus appear to be highly specialized elements of primate upper-layer cortical circuits, raising the question of whether that specialization contributes to their vulnerability.

Beyond our focus on GABAergic interneurons, we also observed an increased abundance of L6b excitatory neurons in SCZ, a finding that initially seemed paradoxical because the adult cortex generates no new neurons. On further consideration, this finding plausibly connects to a large literature describing increased densities of interstitial neurons in the white matter of individuals with SCZ (Akbarian et al. 1993, 1996; Anderson et al. 1996; reviewed in Kubo 2020). Interstitial white matter neurons and L6b neurons are both considered remnants of the developmental subplate (Kostović et al. 2011), and their increased density in SCZ is generally interpreted as neurodevelopmental in origin, reflecting altered migration or failed apoptosis of subplate neurons (Eastwood and Harrison 2003; Duchatel et al. 2019; Kubo 2020). The anatomical source and cause of these excess L6b neurons in SCZ remain unexplored in the present study and warrant dedicated further investigation.

## Limitations

First, observational limits differ by platform; for example, snRNA-seq measures transcripts localized to the nucleus, thus missing mRNAs that preferentially localize to the cytoplasm (Bakken et al. 2018). Likewise, the 300-gene Xenium panel used in our spatial replication analyzes is inherently limited for supertype-level cell typing. Second, all cell typing here was performed using a single reference taxonomy, the SEA-AD MTG neurotypical reference, which we chose because it alone combines supertype-level resolution with spatial transcriptomics and Patch-seq. Third, the depleted cell types we identify are transcriptomically defined, so linking them to morphologically defined types, most notably Sst_25 to the double-bouquet cell, is itself an inference. Fourth, cross-sectional post-mortem sampling cannot temporally order transcriptional from compositional changes, nor separate the effects of disease from those of exposures such as antipsychotic medication. However, the concentration of germline SCZ risk within the depleted Sst supertypes argues that medication exposure alone is unlikely to explain these changes. The donor-level counterpart of this analysis, relating each donor’s polygenic risk to their own cellular composition, would test the link between genetic risk and depletion more directly, but such a test requires genotyped single-nucleus cohorts an order of magnitude larger than any now available. Finally, it would be useful to analyze cellular composition as a function of duration of illness, to determine whether the depleted Sst supertypes are typically depleted before SCZ onset or progressively during the course of the illness.

Excitingly, substantial preclinical work has already sought to develop therapies that compensate for lowered Sst-mediated inhibition, most directly through positive allosteric modulators of α5-subunit-containing GABA-A receptors, which mediate the dendritic inhibition that Sst neurons supply (Fee et al. 2017). Such compounds are advancing towards clinical trials, making them a natural starting point for addressing the deficits described here. However, if Sst cells are indeed being lost in SCZ, as the parallel with AD makes plausible, restoring their inhibitory output will not be enough. Addressing the loss itself requires knowing which cells to protect, and the vulnerable population now has that identity, namely upper-layer Sst supertypes marked by *CALB1* and high *HCN1* expression. The longer-term goal is therefore to understand what renders these neurons vulnerable in the first place, and to protect them before they are lost.

## Acknowledgements

We are deeply grateful to the brain donors and their families, and the investigators and consortia who generated and shared the original datasets analyzed here. We thank Kyle Travaglini for sharing the SEA-AD supertype annotations for the Lee, Dalley et al. Patch-seq cells; Keri Martinowich, Sang Ho Kwon and Cindy Fang for useful discussions and cell annotations related to the Xenium spatial transcriptomics dataset; Yashika Bansal and Netta Ussyshkin for help accessing archival data collected in the Sibille lab; and David Lewis for useful discussions related to somatostatin interneuron diversity and the historical context for their alterations in psychiatric disorders. Large-language models, including Claude, were used in the writing of code and initial drafting of this manuscript.

This work was supported by the Centre for Addiction and Mental Health (CAMH) Discovery Fund, the Krembil Foundation, the Natural Sciences and Engineering Research Council of Canada (NSERC) (RGPIN-2020-05834 and DGECR-2020-00048), the Canadian Institutes of Health Research (CIHR) (PJT-191747, NGN-171423, and PJT-175254), and a Brain Canada Future Leaders in Canadian Brain Research grant. L.D. was supported by the Jaswa Innovator Award from the Stanford Department of Psychiatry, the Uytengsu-Hamilton 22q11 Neuropsychiatry Award from the Stanford Maternal and Child Health Research Institute (MCHRI), and by National Institute of Mental Health (NIMH) grants (nos. R01 MH123486 and R21 MH125358).

## Contributions

N.E. and S.J.T. conceived the study, performed the analyzes, and wrote the manuscript. M.E.F., K.A., X.Z. and T.D. contributed additional analyzes. G.G.-B., L.D. and E.S. contributed additional expertise and guidance that shaped the study and its interpretation. All authors reviewed and approved the final manuscript. S.J.T. supervised the work and acquired funding.

## Data availability

Data generated by this study. Harmonized per-cell cell type annotations for all eight datasets, together with the complete cell type-specific differential expression results at subclass and supertype resolution, are deposited at Zenodo (https://zenodo.org/records/22801230). Supplementary Tables T1–T4 are available with the online version of this paper.

All data analyzed in this study have been previously published and are available in the originating publications and repositories.

Schizophrenia case/control snRNA-seq. Batiuk: Zenodo, https://doi.org/10.5281/zenodo.6921620. McLean and MSSM 1: Synapse, syn25946131 (https://doi.org/10.7303/syn25946131). Multiome: the MultiomeBrain cohort of the PsychENCODE brainSCOPE resource, Synapse syn51111084 (https://doi.org/10.7303/syn51111084; http://brainscope.psychencode.org). Fröhlich: Gene Expression Omnibus, GSE254569. HBCC and MSSM 2: Synapse, syn60084804 (https://doi.org/10.7303/syn60084804).

Spatial transcriptomics. The Xenium DLPFC dataset of Kwon et al. is available from the Gene Expression Omnibus under accession GSE307404.

Reference atlases. The SEA-AD snRNA-seq references (middle temporal gyrus and dorsolateral frontal cortex, A9) and the SEA-AD MTG MERFISH dataset are available from the Seattle Alzheimer’s Disease Brain Cell Atlas (https://portal.brain-map.org/explore/seattle-alzheimers-disease), at https://sea-ad-single-cell-profiling.s3.amazonaws.com and https://sea-ad-spatial-transcriptomics.s3.us-west-2.amazonaws.com respectively.

Human Patch-seq. Intracellular recordings are available from the DANDI Archive (dandiset 000636, https://dandiarchive.org/dandiset/000636); morphological reconstructions and pre-computed electrophysiological features are available from the Allen Brain Map as released with Lee, Dalley et al.

Genetic data. European-ancestry schizophrenia GWAS summary statistics from Bigdeli et al. are available through Synapse (https://www.synapse.org/Synapse:syn60527562); the per-variant export used for the HCN1 locus plot is additionally available from the LocusZoom browser (https://my.locuszoom.org/gwas/282753). Fine-mapped credible sets at the HCN1 locus were taken from Supplementary Table 13 of the same study. PGC3 summary statistics (Trubetskoy et al.) are available from the Psychiatric Genomics Consortium (https://pgc.unc.edu/for-researchers/download-results/).

Gene sets and annotation. MSigDB release 2025.1.Hs (https://www.gsea-msigdb.org/gsea/msigdb), accessed via msigdbr v25.1.1, and org.Hs.eg.db (Bioconductor).

## Code availability

Analysis code is publicly available at https://github.com/stripathy/scz_celltype_paper.

## Ethics and consent

This study consists of secondary analyzes of previously published, de-identified human post-mortem and neurosurgical datasets; tissue collection, donor consent (by donors or next of kin) and ethical approval are described in the source publications cited above. Approval for secondary use of these datasets was provided via CAMH REB #099/2019 titled: Unified proposal for procurement, curation, and secondary analyzes of externally-generated data for mental health research at CAMH.

## Methods

### Data sources and cohorts

#### Schizophrenia case/control single-nucleus RNA sequencing

We assembled seven published post-mortem snRNA-seq datasets of SCZ cases and neurotypical controls, named here by first author or brain bank (Table 1): Batiuk (Batiuk et al. 2022; Zenodo 6921620); HBCC and MSSM 2 (D. Lee et al. 2024; Synapse syn60084804); McLean and MSSM 1 (Ruzicka et al. 2024; Synapse syn25946131); Multiome (Emani et al. 2024; the MultiomeBrain cohort of the PsychENCODE brainSCOPE resource); and Fröhlich (Fröhlich et al. 2024; GEO GSE254569).

Datasets were included if they profiled the prefrontal cortex using 10x Genomics Chromium v3 or v3.1 chemistry and labeled donors as SCZ cases or neurotypical controls. We restricted inclusion to these chemistries because the SEA-AD reference onto which all nuclei were mapped was generated with the same chemistry, and because assigning nuclei to supertypes, rather than subclasses, requires the per-nucleus gene detection that these chemistries provide (Ding et al. 2020). Datasets generated with earlier chemistries, such as the Drop-seq study of Ling et al. (Ling et al. 2024), were therefore not included.

All included datasets sampled prefrontal cortex: dorsolateral prefrontal cortex (DLPFC) for Batiuk, Multiome, HBCC, and MSSM 2; more broadly defined prefrontal cortex (PFC) for McLean and MSSM 1; and orbitofrontal cortex (OFC) for Fröhlich. Where the source publications reported a Brodmann area (BA), these were BA9 (Batiuk), BA10 (McLean), BA9, BA10, and BA46 (MSSM 1), and BA11 (Fröhlich); the Multiome, HBCC, and MSSM 2 datasets are described as sampled from DLPFC with no specific BA designation reported. To limit survivorship bias, only donors younger than 70 years at death were included (cf. Kiss et al. 2026). Furthermore, as the HBCC cohort spans the human lifespan (Yang et al. 2025), this dataset was restricted to donors older than 20 years (range 21–68) to improve age balance between SCZ cases and controls. The meta-analysis comprised 298 controls and 171 SCZ cases across the seven studies.

Together the datasets draw on at least eight brain banks across North America, Europe and Australia (Table 1); Batiuk alone spans four European banks. Two datasets sampled the same bank: MSSM 1 and MSSM 2 both draw on the Mount Sinai NIH NeuroBioBank, and profile matching on sex, diagnosis, age and PMI suggests that 10 donors (8 controls, 2 SCZ cases) may be shared between both datasets. However, as MSSM 1 and MSSM 2 used disparate subject identifiers this overlap could not be confirmed from the identifiers alone, so potentially overlapping donors between datasets were retained as published. Sample sizes, age and sex distributions, brain regions, brain banks and source publications for these and all other re-analyzed datasets are given in Table 1; comprehensive metadata can be found in Supplementary Table T1.

#### Schizophrenia case/control spatial transcriptomics

To test whether snRNA-seq findings replicate in orthogonal, spatially resolved modalities, we additionally analyzed a Xenium dataset of post-mortem human DLPFC (BA46) from the Lieber Institute for Brain Development (LIBD) Human Brain and Tissue Repository, comprising 12 SCZ cases and 12 neurotypical controls (Kwon et al. 2026). These data were obtained from Gene Expression Omnibus (GEO GSE307404).

#### SCZ GWAS summary statistics

Cell-type genetic enrichment used the European-ancestry autosomal meta-analysis of Bigdeli et al. 2026 (Bigdeli et al. 2026) (GRCh38; maximum per-variant effective N = 114,827). Fine-mapped credible sets at the *HCN1* locus were taken from the same study’s SuSiE-R European-ancestry analysis. The PGC3 meta-analysis (Trubetskoy et al. 2022; European-ancestry autosomal) was used for robustness comparisons.

#### SEA-AD reference atlases

Four components of the Seattle Alzheimer’s Disease Brain Cell Atlas (SEA-AD; Gabitto et al. 2024) were used. (1) The MTG snRNA-seq neurotypical reference (five donors, 137,303 nuclei) defines the 24-subclass and 137-supertype taxonomy onto which all cell typing in this study is harmonized, including snRNA-seq label transfer and Xenium cell typing. (2) The dorsolateral frontal cortex (A9) snRNA-seq release provided region-matched normotypic expression for the genetic-enrichment analyzes, restricted to its three neurotypical reference donors (90,579 nuclei; 125 supertypes carrying the canonical labels). (3) The aged SEA-AD cohorts provided the Alzheimer’s disease comparison, where each donor carries a pathology-based continuous pseudo-progression score (CPS) indexing AD neuropathological severity based on sections from MTG, and the DLPFC cohort (80 donors with CPS and complete covariates) was used for the primary AD compositional axis, with the MTG cohort used to test regional robustness. (4) The MTG MERFISH dataset (27 donors) was used for the Xenium cortical-depth analyzes — training and validating the depth model — and for benchmarking Xenium cell-type proportions.

#### Human Patch-seq recordings

Electrophysiology and morphology came from the human cortical patch-seq dataset of Lee, Dalley et al. 2023 (B. R. Lee et al. 2023): 778 unique GABAergic interneurons from 152 donors, recorded in acute and cultured slices prepared from neurosurgical resections (epilepsy, tumor, or vascular malformation). Sampled areas follow surgical access rather than a sampling design, and are predominantly temporal but include prefrontal and cingulate cortex. Pre-calculated intrinsic electrophysiological features and biocytin morphological reconstructions were used as released in the source publication.

### Transcriptomics data processing and cell-type harmonization

#### snRNA-seq

For each of the seven snRNA-seq datasets we started from the authors’ quality-controlled, filtered count matrices. Because each study had annotated its nuclei with a different taxonomy, we re-annotated every nucleus against a single reference, the SEA-AD MTG taxonomy of 24 subclasses and 137 supertypes defined from five neurotypical donors (described above), using Seurat reference-based label transfer (Stuart et al. 2019). Label transfer was run separately for each dataset. Reference and query were restricted to their shared genes and log-normalized (scale factor 1,000,000); the 3,000 most variable genes were selected by variance-stabilizing transformation; the reference was scaled and reduced by principal component analysis; transfer anchors were identified in the first 30 principal components; and labels with their prediction scores were assigned with Seurat’s TransferData. Each nucleus thus received a class, subclass (e.g., Sst) and supertype (e.g., Sst_25) label that is used consistently across datasets and analyzes, and all downstream analyzes used the original, unnormalized counts. To check the transferred labels, we compared canonical marker-gene expression between the seven datasets combined and the SEA-AD reference at subclass and Sst-supertype resolution, which showed the expected marker profiles (Fig. S1).

Two datasets (HBCC and MSSM 2) were too large to hold in memory at once and were therefore split at random into subsets that were label-transferred independently; this changes nothing about the procedure applied to any individual nucleus. For the same two datasets, Ensembl gene identifiers were converted to gene symbols with org.Hs.eg.db (Carlson et al. 2019).

For the purposes of visualization, UMAP embeddings of all datasets together (Fig. 1b–d) were computed with brisc (https://github.com/briscverse/brisc), which is designed to handle large datasets of this size. Cell-type marker genes were identified using brisc’s find_markers function. Top markers were selected based on ranked positive logFC and the top three markers for each cell type were selected for visualization.

#### Xenium

We reanalyzed the Xenium DLPFC dataset of Kwon et al. (2026); 24 sections, 12 SCZ / 12 control; 1.34 million cells; 300-gene panel). Because 300 genes are too few for de novo clustering, each cell was assigned to an SEA-AD subclass and supertype by a two-stage correlation classifier built on the same reference, and to a continuous cortical depth (0 = pia, 1 = white matter) by a neighborhood-composition model; both were validated against the independent SEA-AD MTG MERFISH atlas, and the supertype resolution attainable with the panel was characterized by a leave-one-donor-out benchmark on the snRNA-seq reference (Figs. S2–S3). All disease analyzes used one shared cell set, comprising 742,103 total cells (356,313 neuronal, 385,790 non-neuronal) across the 24 sections that passed spatial, cell type labeling, and doublet-based quality control and with cell soma within cortical layers 1–6 (i.e., not annotated to vasculature/meninges or to white matter); their differential expression and composition analyzes are described in the corresponding sections below. Comprehensive cell annotations for the snRNA-seq datasets and the Xenium dataset are deposited at Zenodo (https://zenodo.org/records/22801230).

### Cell type-specific differential gene expression

#### snRNA-seq

To quantify how SCZ alters gene expression within cell types, we tested differential expression (DE) separately in each of the seven snRNA-seq datasets at SEA-AD subclass and supertype resolution. Within each dataset, nuclei were aggregated into per-donor, per-cell-type pseudobulk profiles using Seurat (Hao et al. 2024). To ensure robust estimation, for subclasses, only donors contributing ≥ 500 nuclei across all subclasses were retained. For the supertype level, only donors contributing ≥ 10 cells of the given cell type were retained. For both subclass and supertype analyzes only genes with ≥ 1 count in at least 80% of retained donors were tested. Counts were TMM-normalized (edgeR; Robinson et al. 2010), transformed with voom, and tested with limma’s moderated t-statistics (Law et al. 2014; Ritchie et al. 2015). SCZ diagnosis was the predictor of interest, with age at death, sex and PMI as covariates, age and PMI standardized (1); PMI was unavailable for the Multiome dataset and was omitted from that dataset’s model.

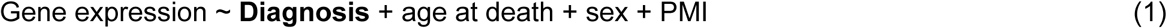

#### Meta-analysis

Per-dataset log_2_ fold changes and standard errors were obtained for each gene × cell-type pair, and pairs with estimates from at least five datasets were pooled by random-effects meta-analysis (metafor::rma, REML;Viechtbauer 2010), which weights each dataset by the inverse of its within-dataset variance plus the estimated between-dataset variance and so allows the effect of SCZ on a gene to differ between datasets. Pooled p-values were Benjamini–Hochberg false discovery rate (BH-FDR)-adjusted independently within each cell type. Low-abundance supertypes were not meta-analyzed when fewer than five datasets had sufficient cell counts to retain enough donors for differential expression analysis.

#### Xenium

For spatial replication, we applied the same design to the Xenium dataset. Counts were summed into per-section, per-subclass pseudobulk profiles from cell types with ≥ 10 cells in a section, and genes detected in ≥ 80% of retained sections were tested with an edgeR (Robinson et al. 2010) quasi-likelihood F-test, chosen over voom-limma because the 300-gene panel gives too few genes to estimate voom’s mean–variance trend reliably. DE was tested under the same model (1, as above) and BH-FDR-corrected within each subclass across the 300-gene panel. Because this FDR is computed over far fewer genes, it is not directly comparable to the genome-wide snRNA-seq FDR.

#### Consistency of DE between snRNA-seq and Xenium

To assess cross-platform concordance we intersected meta-analytic DE gene × subclass pairs (meta FDR < 0.10) with the pairs testable in Xenium, correlated the two platforms’ log₂ fold changes and tested directional agreement with a binomial sign test.

#### Gene-set enrichment analyses

To compare transcriptional dysregulation between Sst supertypes that are depleted and those that persist, nuclei were pooled across the supertypes of each depletion group (depleted, intermediate, non-depleted; defined in Results) into one pseudobulk profile per donor per dataset, and DE and meta-analysis were run as above, with BH-FDR correction within each group. Gene-set enrichment analysis was then performed with fgsea (Korotkevich et al. 2021) on genes ranked by the meta-analytic z statistic (estimate/standard error), separately for each group and for the pooled Sst subclass. Gene sets used comprised the Gene Ontology biological process, cellular component and molecular function collections and Reactome, from MSigDB release 2025.1.Hs (Liberzon et al. 2015) via msigdbr v25.1.1, restricted to 10–500 genes (6,024–6,839 sets per group) and tested with 10,000 permutations; significance was assessed at BH-FDR < 0.10 within each group. Leading-edge genes are fgsea’s leading-edge subset, the ranked genes contributing to a set’s enrichment score.

### Cell type-specific differential abundance analyzes

#### snRNA-seq

To test whether SCZ alters the relative abundance of cell types, we modeled cell-type composition in each of the seven snRNA-seq datasets at supertype resolution using crumblr (Hoffman and Roussos 2025). Crumblr applies a centered-log-ratio transform to each donor’s cell-type counts and fits precision-weighted linear models, so that donors contributing more nuclei, whose proportions are estimated more precisely, carry more weight. Because the neuronal fraction of nuclei varies widely between samples and datasets for technical reasons, above all differences in gray- versus white-matter content, we followed SEA-AD (Gabitto et al. 2024) in analyzing neuronal and non-neuronal cells as two separate compositions. Every proportion reported here is therefore within class, for example the fraction of all neurons assigned to a given neuronal supertype. Within each dataset, nuclei were counted per donor and supertype, and a supertype was modeled in a dataset if at least one nucleus in that dataset was assigned to it. Each model had SCZ diagnosis as the predictor of interest and age at death, sex and PMI as covariates, with age and PMI standardized (2); PMI was unavailable for the Multiome dataset and was omitted from that dataset’s model. Each beta coefficient was tested with a moderated t-statistic, in which the residual variance of each cell type is shrunk toward the variance pooled across cell types in that dataset (limma’s empirical Bayes procedure), and P values were BH-FDR-corrected across cell types within each dataset.

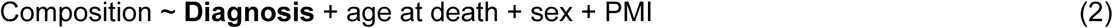

#### Meta-analysis

Per-dataset coefficients and standard errors were pooled across the seven datasets by fixed-effect meta-analysis (metafor::rma, FE; Viechtbauer 2010), which weights each dataset by the inverse of its variance, and pooled p-values were BH-FDR-adjusted across supertypes within each class. A supertype was meta-analyzed if at least two datasets provided a valid estimate. We chose fixed-effect pooling as our primary method for pooling estimates because the seven datasets were analyzed with one taxonomy and one model, and several datasets are somewhat small (11–35 donors), so the between-dataset variance on which random-effects weights depend is poorly estimated.

#### Sensitivity to meta-analysis pooling strategy and to single datasets

We repeated the dataset pooling using multiple alternative strategies to assess sensitivity to this choice. First, the per-dataset estimates were combined by random-effects meta-analysis (metafor::rma, REML), which also yields the between-dataset heterogeneity of each supertype (I² and Cochran’s Q). Second, the seven datasets were analyzed jointly in a mega-analysis that fit all donors across datasets in a single crumblr model using variancePartition::dream (Hoffman and Roussos 2021), with dataset as a random intercept, (1 | dataset), or as a random intercept together with a dataset-specific random SCZ slope, (1 + Diagnosis | dataset). Intuitively, the first allows each dataset its own baseline composition while estimating one SCZ effect from all donors, the donor-level counterpart of fixed-effect pooling; the second additionally lets the SCZ effect vary between datasets around a common mean, so that the pooled estimate and its uncertainty reflect between-dataset disagreement, the counterpart of random-effects pooling. Here, age and PMI were standardized across all donors, and PMI values that were unavailable (Multiome) or recorded as 0 h (one MSSM 2 donor) were set to the mean. Lastly, to assess dependence of effects on any single dataset, the fixed-effect meta-analysis was repeated with each dataset omitted in turn, recomputing BH-FDR within each re-analysis.

#### Sensitivity to DE genes in label transfer

Because differential expression in SCZ could bias cell type label transfer, a shift in composition could in principle reflect mislabeling rather than a change in cell number. To test this, we removed the genes that were DE in Sst cells at the subclass level at FDR < 0.10 from the count matrices of all seven datasets and of the reference, re-annotated every nucleus with the label-transfer method described above, and re-ran the crumblr pipeline on the resulting counts.

#### Xenium

For spatial replication, we applied the same design to the Xenium dataset (Kwon et al. 2026; cell set as defined above). Cells were counted per section (one section per donor; n = 24) and per cell type, using cortical cells only (those annotated to layers 1–6); neuronal and non-neuronal classes were again modeled separately with within-class totals as the denominator, and the model of (2) was fit at subclass and at supertype resolution. BH-FDR correction was performed across cell types within each class at each level.

### Consistency of composition between snRNA-seq and Xenium

To assess cross-platform concordance, we correlated the SCZ beta coefficients of the snRNA-seq meta-analysis with those of Xenium across all cell types analyzed on both platforms.

#### Alzheimer’s disease pseudo-progression

To ask whether the supertypes depleted in SCZ also decline in AD, we applied the same crumblr model to the 80 donors of the SEA-AD DLPFC aged dataset (neurotypical reference donors excluded), replacing the diagnosis term with each donor’s continuous pseudo-progression score (CPS):

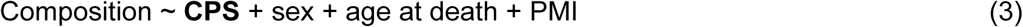

with CPS, age and PMI standardized and each donor’s total neuronal nuclei as the compositional denominator. The coefficient of interest is the compositional slope per standard deviation of CPS; negative values mark supertypes that decline as AD advances. Because SEA-AD uses the same supertype taxonomy as the rest of this study, these slopes are directly comparable to the SCZ coefficients. As a regional check, the identical model was fit to the SEA-AD MTG aged dataset, the same 80 donors with only the dissected region differing.

### Cell-type enrichment of schizophrenia genetic risk

#### Overview

Linking common-variant genetic risk to cell types requires two inputs: a GWAS and a reference expression atlas for cell types. The approach of Duncan et al. 2025 (Duncan et al. 2025) reduces each to a per-gene quantity: 1) a gene-level association statistic from the GWAS, and 2) a cell type-specific expression specificity score from the atlas measuring how concentrated a gene’s expression is in one cell type relative to all others. The method asks, for each cell type in turn, whether the genes expressed most specifically in that type are also those carrying the most genetic association. We followed this approach, updating two of its inputs: the SCZ GWAS (Bigdeli et al. 2026), and the expression atlas, for which we used the SEA-AD supertypes in place of the Siletti whole-brain clusters, so that the cell types carrying genetic signal are the same supertypes whose abundance we measured in disease.

#### Expression reference

We used the SEA-AD DLPFC (A9) reference as the primary resource for deriving normotypic expression levels of SEA-AD supertypes, because expression specificity is quantitative and may be sensitive to regional differences in expression levels, and DLPFC matches the prefrontal origin of the SCZ case-control data used here. Because specificity is computed across the 125 supertypes present in DLPFC (the 12 absent are non-neuronal or vascular apart from L5 ET_1; all 16 Sst supertypes are present), each gene’s value for a supertype is its share of expression relative to the other cortical types rather than to a brain-wide set. For robustness, the same analysis was repeated with specificity computed from the MTG reference.

#### Specificity and enrichment

Cell type-specific expression specificity was computed as specified by Duncan et al. (Duncan et al. 2025): per-cell ln(1 + raw UMI) averaged within type; genes restricted to unique symbols and unambiguous one-to-one mappings between Entrez and Ensembl identifiers (org.Hs.eg.db), with the extended MHC excluded; each cell type scaled to a common total; and each gene then normalized across types, so that its value for a type is that type’s share of the gene’s total expression (16,544 genes × 125 types). Gene-level association used MAGMA v1.10 (de Leeuw et al. 2015) with a 35 kb upstream / 10 kb downstream annotation window and the SNP-wise mean model against the 1000 Genomes phase 3 European panel, with per-variant effective N. The LD panel and gene coordinates are hg19, as distributed with MAGMA, and must share a build with one another; summary statistics were therefore joined to the panel by rsID and the panel’s own positions used throughout. Cell-type association was tested with MAGMA’s gene-property analysis (one-sided, positive) at its defaults, over the 15,981 genes common to both inputs. Each supertype is tested in its own regression, so the 125 supertypes do not compete with one another. P values were Bonferroni- and BH-FDR-corrected across the 125 tested supertypes.

#### Gene drivers

For Sst_2, driver genes were defined as genes jointly in the top 10% of Sst_2 specificity and passing MAGMA gene-level FDR < 0.05 among the 15,855 tested genes (344 genes).

### Molecular characterization of the depleted Sst supertypes in normotypic reference data

Sst supertypes were assigned to depleted and not-depleted groups from the compositional snRNA-seq meta-analysis: the five supertypes reduced in SCZ at meta-analytic FDR < 0.20 (Sst_2, Sst_3, Sst_20, Sst_22 and Sst_25) formed the depleted group, and the remaining eleven Sst supertypes the not-depleted group. Expression levels from the two groups were then compared in the SEA-AD DLPFC A9 neurotypical reference. Counts were normalized to 10,000 per nucleus and log1p-transformed, genes tested by cell-level Wilcoxon rank-sum with BH correction, and Seurat-style avg_log_2_FC computed on the un-logged CP10K scale; genes were called differentially expressed at FDR < 0.05 and |log_2_FC| > 0.25.

### Patch-seq electrophysiology and morphology

#### Supertype assignment

To link the SCZ-depleted Sst supertypes to their physiological and morphological properties, we used SEA-AD supertype calls for the Lee, Dalley et al. Patch-seq cells (Data sources). These are the same scANVI-derived supertype assignments, for the same cells, that were used in the SEA-AD study (Gabitto et al. 2024), provided directly by K. Travaglini (personal communication). Of the 757 Lee/Dalley Patch-seq characterized cells receiving a label, 156 were labeled as belonging to the Sst subclass; we used the 150 cells that also carried a recorded cortical layer, spanning 16 Sst supertypes collected from 59 distinct donors.

#### Physiology and depth

Per-supertype mean sag ratio was computed over the 129 cells carrying an extracted sag value, using the released feature values (Data sources). Sag ratio was calculated as the fraction of the peak hyperpolarizing voltage deflection that relaxes back toward baseline by the end of the current step, (V_peak − V_steady)/(V_peak − V_baseline), measured on the step whose peak deflection is closest to −100 mV; larger values indicate stronger sag, the signature of HCN-channel-mediated I_h. Five biocytin-recovered reconstructions spanning the range of soma depths are shown soma-aligned to pia and ordered by depth; soma depth was measured on each reconstruction as the distance from pia, and voltage responses were read from the corresponding electrophysiological sweeps.

