## Supplementary Figures for "Selective depletion of upper-layer somatostatin interneuron subtypes in schizophrenia"

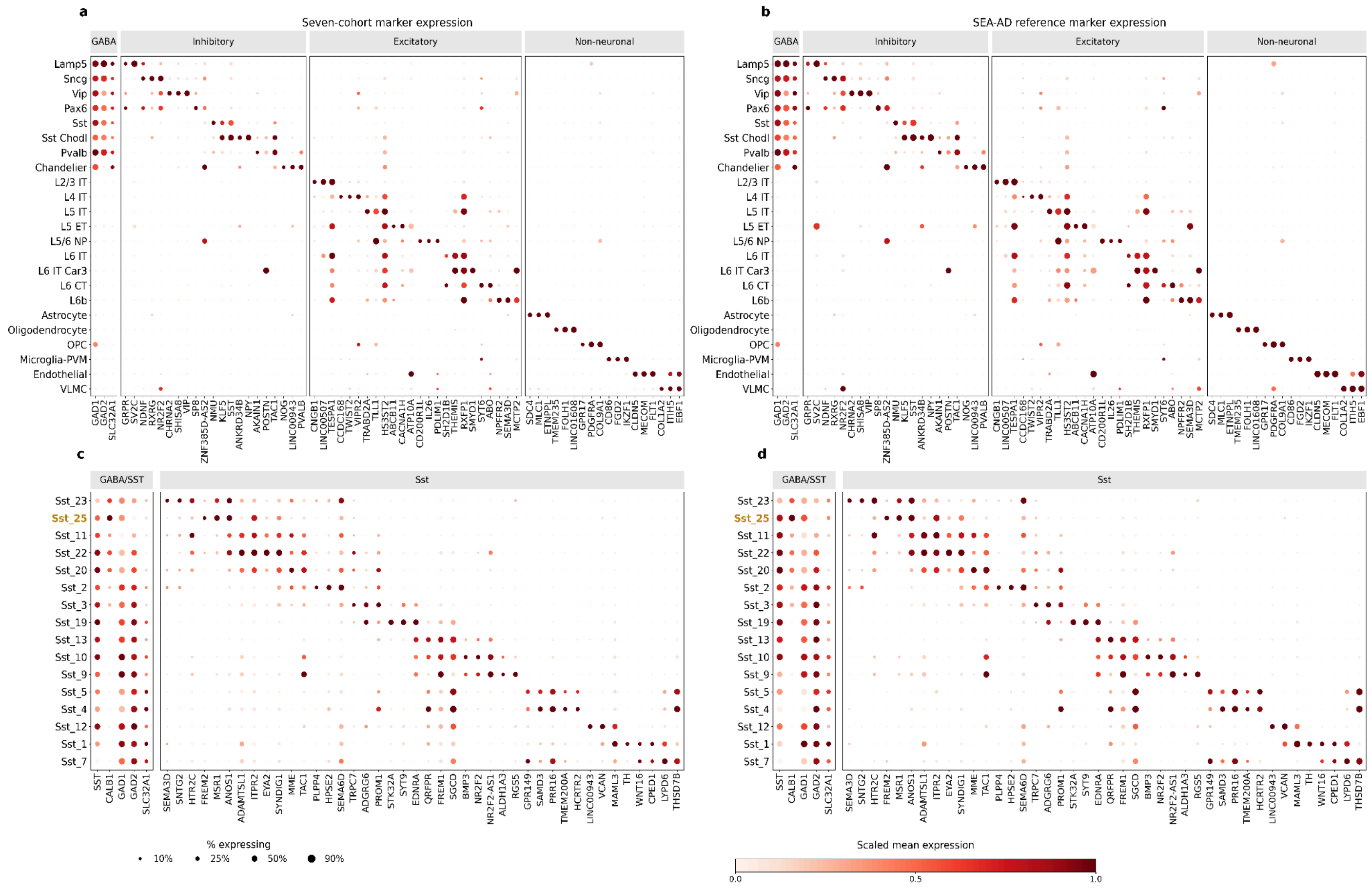


**Fig. S1 | Marker dot plots showing canonical marker expression across predicted cell types.**

Marker expression is shown for the seven-cohort integrated dataset (**a**,**c**) and the SEA-AD reference (**b**,**d**). Subclass-level panels (**a**,**b**) are grouped into inhibitory, excitatory, and non-neuronal cell classes, whereas Sst-supertype panels (**c**,**d**) are ordered approximately from pia to white matter. Dot size indicates the fraction of cells expressing each gene, and colour indicates scaled mean expression.

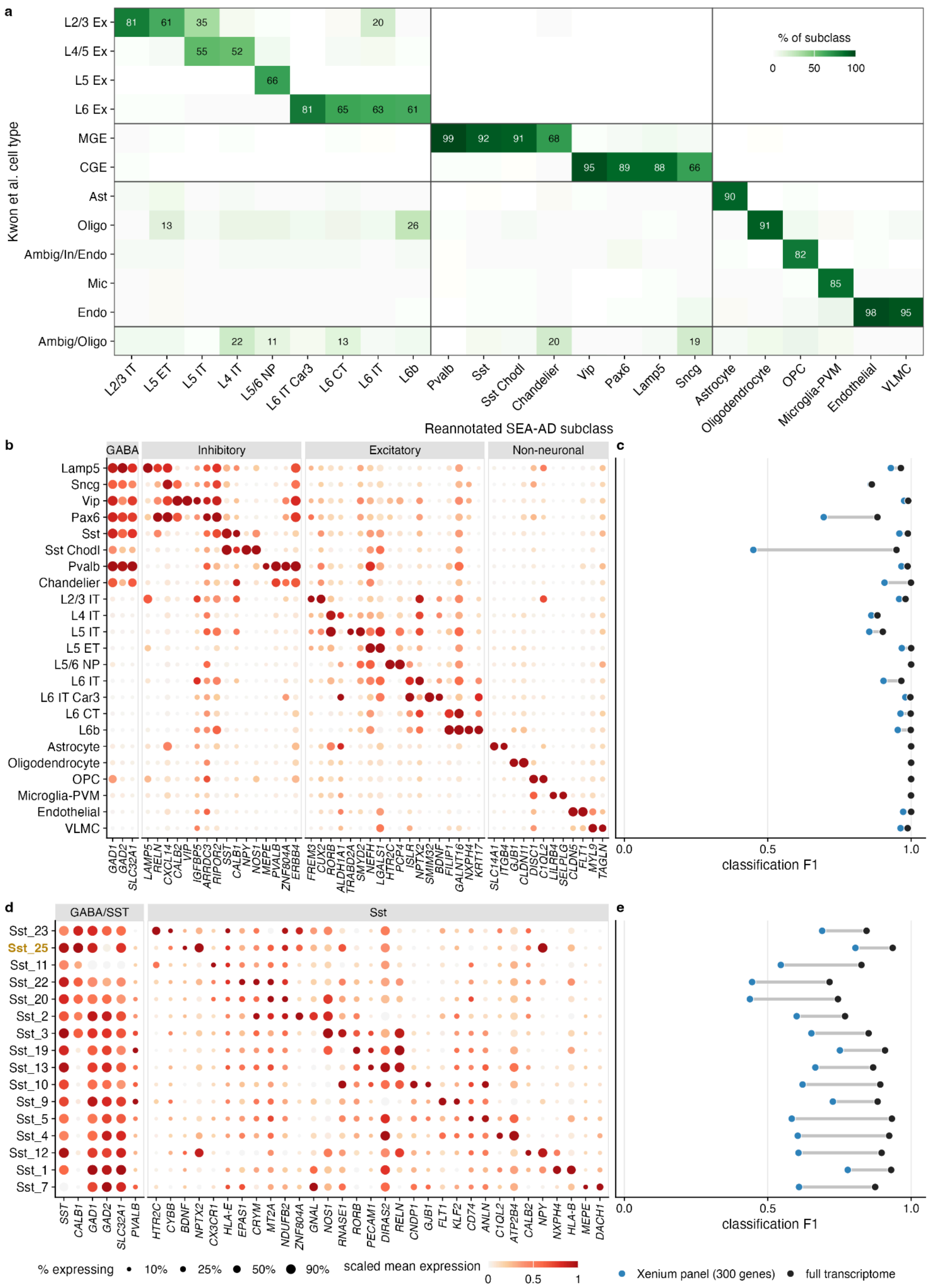
a
% of subclass
0
50
100
Kwon et al. cell type
L2/3 Ex
L4/5 Ex
L5 Ex
L6 Ex
MGE
CGE
Ast
Oligo
Ambig/In/Endo
Mic
Endo
Ambig/Oligo
81
61
35
20
55
52
66
81
65
63
61
99
92
91
68
95
89
88
66
90
13
26
91
82
85
98
95
22
11
13
20
19
L2/3 IT
L5 ET
L5 IT
L4 IT
L5/6 NP
L6 IT Car3
L6 CT
L6 IT
L6b
Pvalb
Sst
Sst Chodl
Chandelier
Vip
Pax6
Lamp5
Sncg
Astrocyte
Oligodendrocyte
OPC
Microglia-PVM
Endothelial
VLMC
Reannotated SEA-AD subclass
b
GABA
Inhibitory
Excitatory
Non-neuronal
Lamp5
Sncg
Vip
Pax6
Sst
Sst Chodl
Pvalb
Chandelier
L2/3 IT
L4 IT
L5 IT
L5 ET
L5/6 NP
L6 IT
L6 IT Car3
L6 CT
L6b
Astrocyte
Oligodendrocyte
OPC
Microglia-PVM
Endothelial
VLMC
GAD1
GAD2
SLC32A1
LAMP5
RELN
CXCL14
CALB2
VIP
IGFBP5
ARRDC3
RIPOR2
SST
CALB1
NPY
NOS1
MEPE
PVALB
ZNF804A
ERBB4
FREM3
CUX2
RORB
ALDH1A1
TRABD2A
SMYD2
NEFH
LGALS1
HTR2C
PCP4
ISLR
NPTX2
SMIM32
BDNF
FILIP1
GALNT16
NXPH4
KRT17
SLC14A1
ITGB4
GJB1
CLDN11
DISC1
C1QL2
LILRB4
SELPLG
CLDN5
FLT1
MYL9
TAGLN
c
0.0
0.5
1.0
classification F1
d
GABA/SST
Sst
Sst_23
Sst_25
Sst_11
Sst_22
Sst_20
Sst_2
Sst_3
Sst_19
Sst_13
Sst_10
Sst_9
Sst_5
Sst_4
Sst_12
Sst_1
Sst_7
SST
CALB1
GAD1
GAD2
SLC32A1
PVALB
HTR2C
CYBB
BDNF
NPTX2
CX3CR1
HLA-E
EPAS1
CRYM
MT2A
NDUFB2
ZNF804A
GNAL
NOS1
RNASE1
RORB
PECAM1
DIRAS2
RELN
CNDP1
GJB1
FLT1
KLF2
CD74
ANLN
C1QL2
ATP2B4
CALB2
NPY
NXPH4
HLA-B
MEPE
DACH1
e
0.0
0.5
1.0
classification F1
% expressing
10%
25%
50%
90%
scaled mean expression
0
0.5
1
Xenium panel (300 genes)
full transcriptome

**Fig. S2 | Xenium cell-type annotation illustrating agreement with the original annotations, panel marker genes, and cell type resolvability given gene panel.**

(**a**) Reannotated SEA-AD subclass calls against the dataset authors' (Kwon et al.) independent annotations; each cell is the percentage of a reannotated subclass assigned to a given author cell type (columns sum to 100), values ≥ 10% labeled, ordered to place the dominant correspondence on the diagonal. (**b, d**) Marker dot-plots at subclass level (b; inhibitory, excitatory, non-neuronal) and Sst-supertype level (d; ordered pia to white matter); dot size, fraction of cells expressing; fill, scaled mean expression. (**c, e**) Classification F1 on the 300-gene Xenium panel (blue) versus the full transcriptome (black) using a nearest-centroid Pearson classifier under leave-one-donor-out cross-validation on the SEA-AD neurotypical snRNA-seq reference at the subclass (**c**) and Sst supertype (**e**) level. Rows in b/c and d/e share the same cell type y-axis labels shown at left.

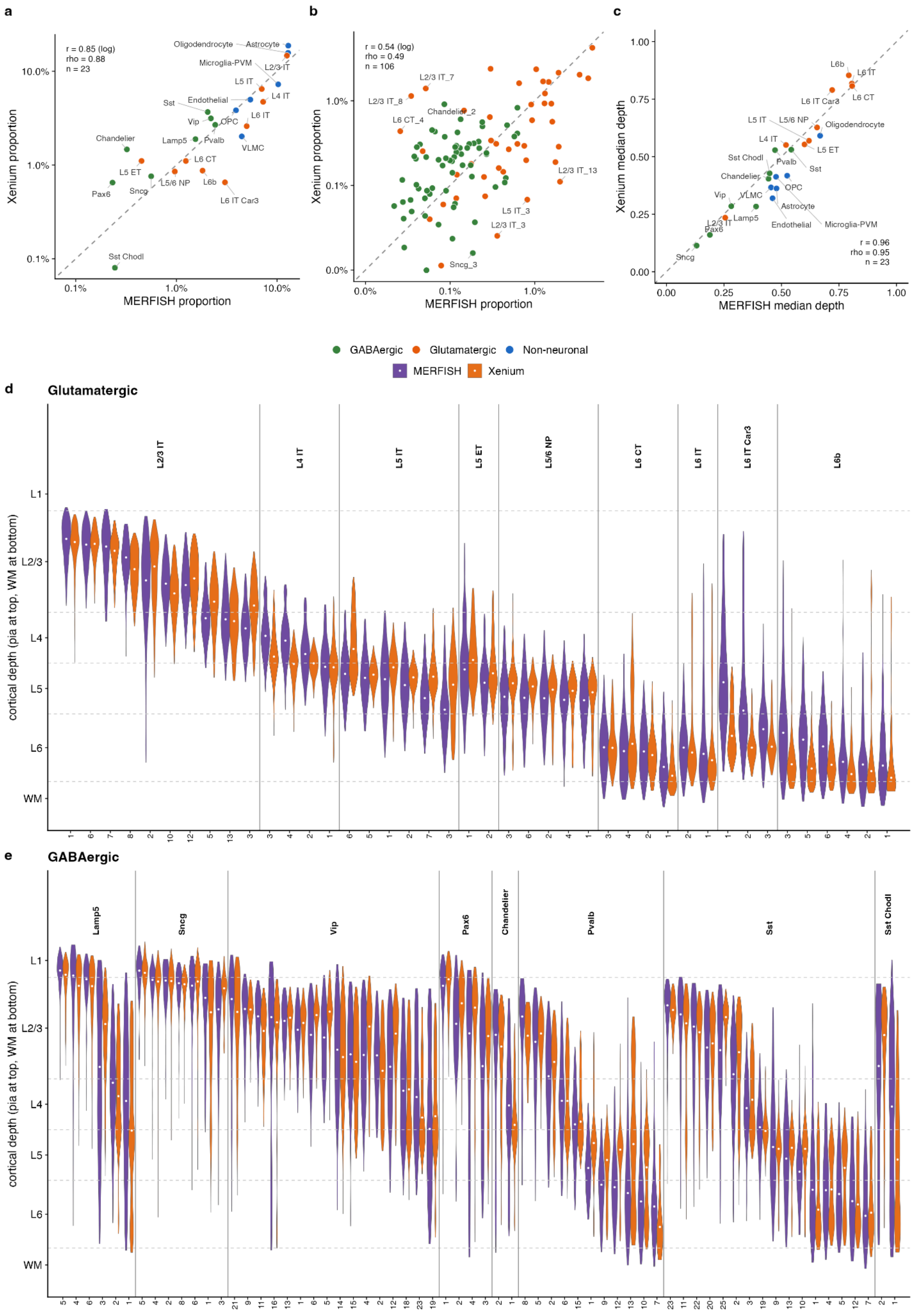
a
r = 0.85 (log)
rho = 0.88
n = 23
Xenium proportion
MERFISH proportion
Oligodendrocyte
Astrocyte
Microglia-PVM
L2/3 IT
L5 IT
L4 IT
Endothelial
Sst
L6 IT
Vip
OPC
Chandelier
Lamp5
Pvalb
VLMC
L6 CT
L5 ET
L5/6 NP
L6b
Sncg
Pax6
L6 IT Car3
Sst Chodl
b
r = 0.54 (log)
rho = 0.49
n = 106
L2/3 IT_7
L2/3 IT_8
Chandelier_2
L6 CT_4
L2/3 IT_13
L5 IT_3
L2/3 IT_3
Sncg_3
c
Xenium median depth
MERFISH median depth
r = 0.96
rho = 0.95
n = 23
L6b
L6 IT
L6 CT
L6 IT Car3
L5 IT
L5/6 NP
Oligodendrocyte
L4 IT
L5 ET
Sst Chodl
Pvalb
Sst
Chandelier
OPC
Vip
VLMC
Astrocyte
L2/3 IT
Pax6
Lamp5
Endothelial
Microglia-PVM
Sncg
GABAergic
Glutamatergic
Non-neuronal
MERFISH
Xenium
d
Glutamatergic
cortical depth (pia at top, WM at bottom)
L1
L2/3
L4
L5
L6
WM
L2/3 IT
L4 IT
L5 IT
L5 ET
L5/6 NP
L6 CT
L6 IT
L6 IT Car3
L6b
e
GABAergic
Lamp5
Sncg
Vip
Pax6
Chandelier
Pvalb
Sst
Sst Chodl

**Fig. S3 | Xenium vs SEA-AD MERFISH concordance: cell-type proportions and cortical depth.**

(**a**) Per-donor subclass proportions, Xenium vs MERFISH (Pearson r = 0.85, $\log_{10}$; ρ = 0.88, n = 23). (**b**) Neuronal supertype proportions (r = 0.54, $\log_{10}$; ρ = 0.49, n = 106), illustrating the panel's within-subclass limit. (**c**) Subclass median cortical depth (r = 0.96; ρ = 0.95, n = 23). (**d, e**) Per-supertype cortical depth distributions (0 = pia, 1 = WM) in MERFISH (purple; manual annotation) and Xenium (orange; model prediction), for the 42 glutamatergic (**d**) and 64 GABAergic (**e**) supertypes of b; grouped by subclass and ordered pia → WM by MERFISH median; violins show the inner 95% of cells, white point = median; dashed lines, laminar boundaries. Median depths agree at r = 0.97 across these supertypes and at r = 0.94 within subclass, though Xenium distributions are systematically narrower (median IQR 0.120 vs 0.145).

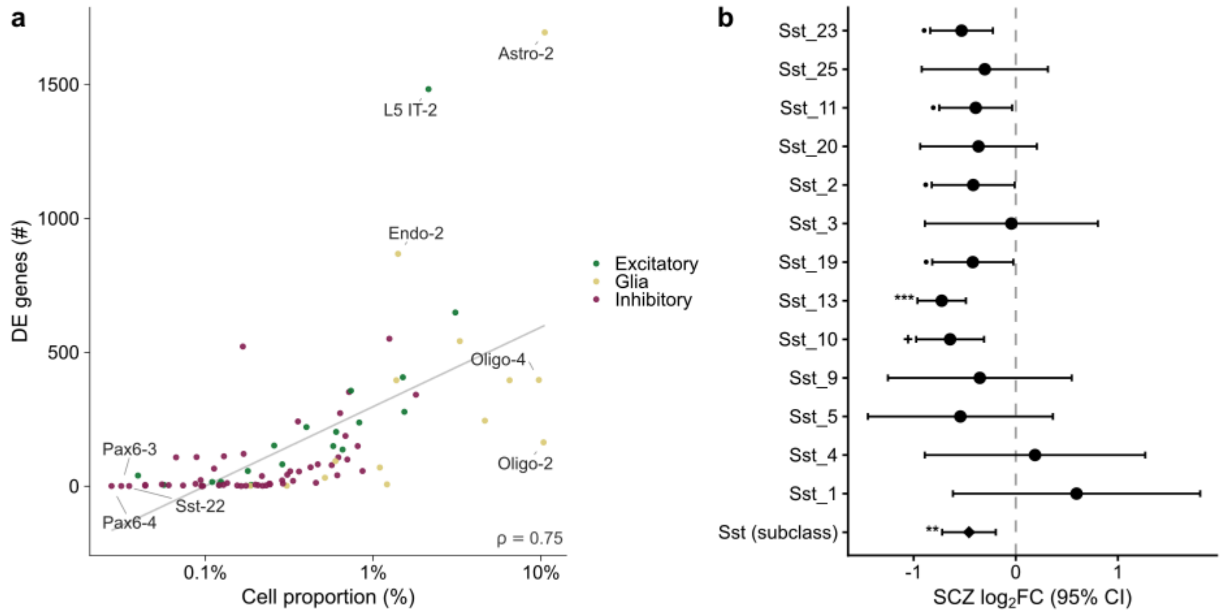


**Fig. S4 | Supertype-level differential-expression**

(**a**) Scatter plot showing the relationship between mean cell-type abundance and the number of differentially expressed (DE) genes for each supertype. Cell abundance was calculated as the mean proportion of total nuclei contributed by each supertype across donors. DE gene counts represent the number of unique genes significant at FDR < 0.10 across the schizophrenia snRNA-seq meta-analysis. Each point represents a supertype and is colored according to its cell class. The x-axis is shown on a log10 scale. The gray line indicates the linear trend, and ρ denotes the Spearman rank correlation between cell-type abundance and the number of DE genes. (**b**) Forest plot showing SCZ-associated log2FC in the expression of SST mRNA among cell types. Points indicate effect estimates and horizontal bars indicate 95% confidence intervals; the diamond denotes the Sst subclass-level estimate. The dashed vertical line indicates log2FC = 0. Significance markers indicate *** FDR < 0.01, ** FDR < 0.05, * FDR < 0.10, + FDR 0.10–0.20, and • unadjusted P < 0.05.

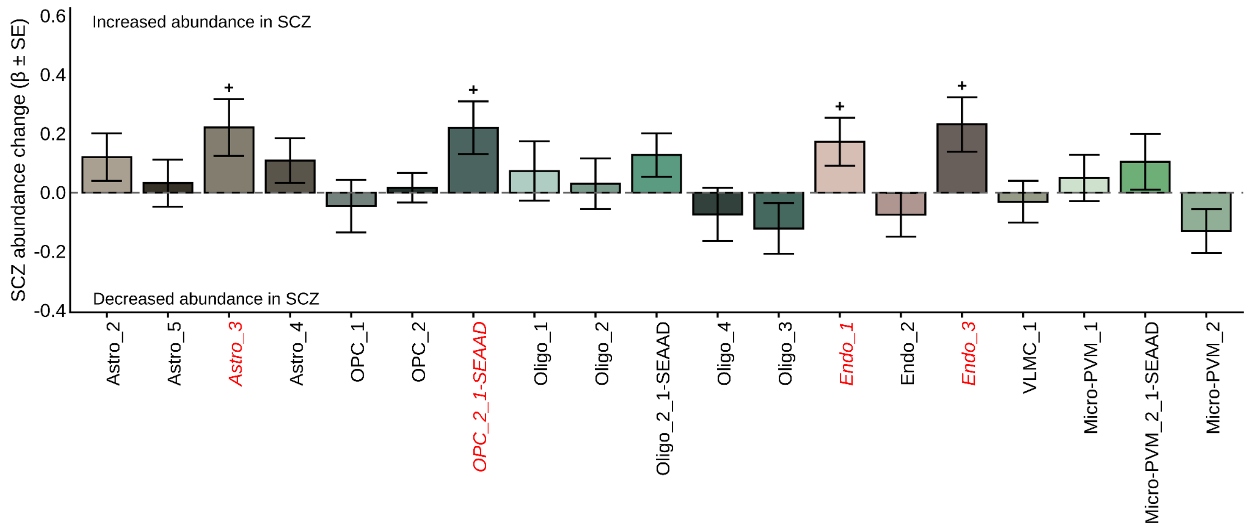


**Fig. S5 | Non-neuronal abundance changes in schizophrenia in snRNA-seq.**

Cell type-specific abundance changes in schizophrenia (crumblr β ± SE) for each of the 19 non-neuronal supertypes meta-analyzed across seven snRNA-seq datasets (N = 298 control / 171 SCZ donors), covarying age, sex and post-mortem interval. Positive bars denote increased cell type abundance in SCZ; negative bars denote decreased or depleted cell types. Bars are colored by supertype. Supertype labels in italic red indicate trend-level changes (FDR < 0.20); markers above or below bars denote statistical significance: + FDR 0.10–0.20.

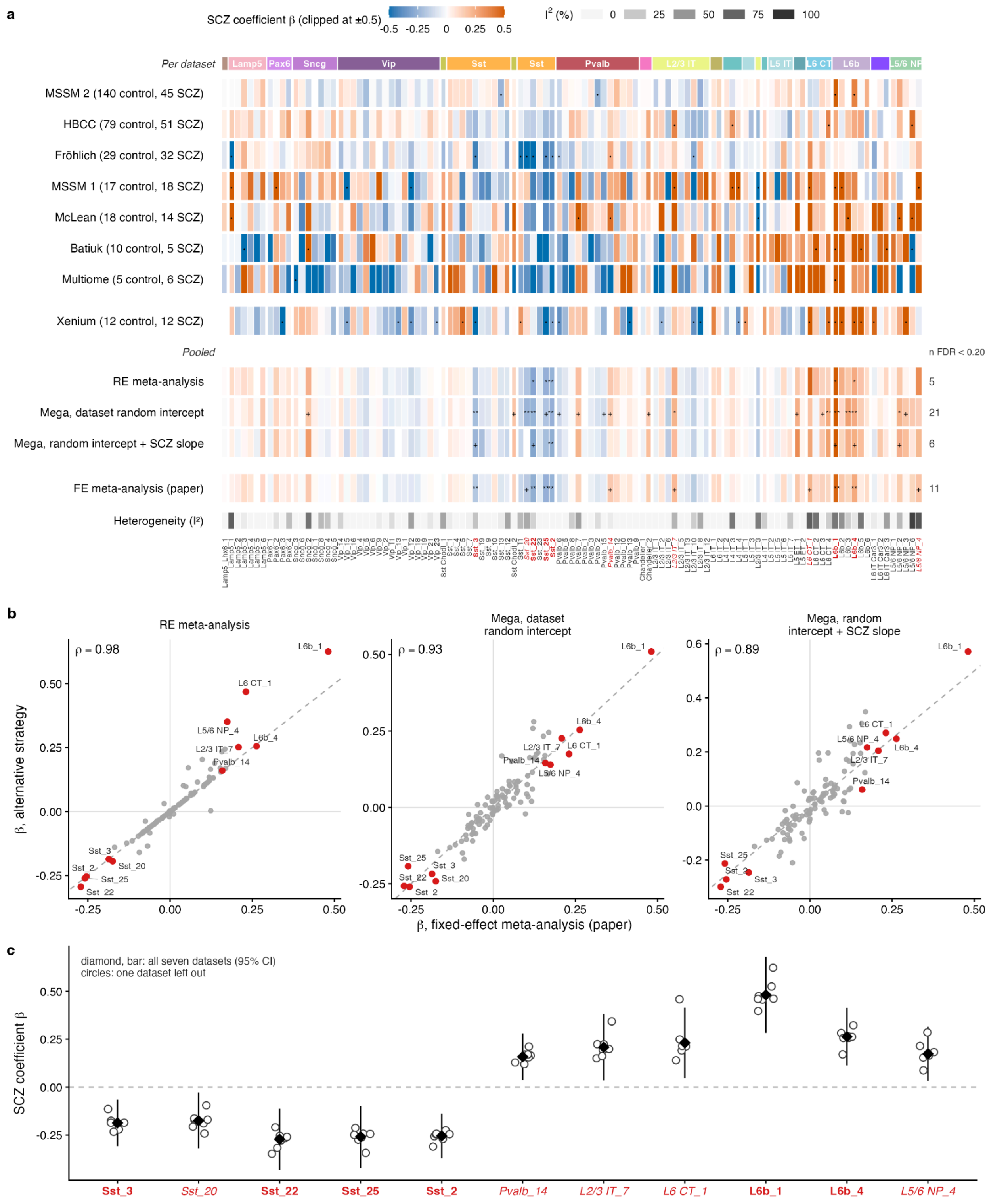


**Fig. S6 | Robustness of the snRNA-seq cell-type abundance results to individual datasets and to the pooling strategy.**

(**a**) SCZ abundance change (crumblr β; red, higher relative abundance in SCZ; blue, lower; clipped at ±0.5) for the 109 neuronal supertypes, grouped by subclass (colored strip). Top block, each dataset analyzed on its own: the seven snRNA-seq datasets and the Xenium dataset, with control and SCZ donor numbers in parentheses; dot, nominal $P < 0.05$. Lower block, various estimates pooled across the seven snRNA-seq datasets: a random-effects meta-analysis of the per-dataset estimates; two mega-analyzes that fit all donors across datasets in a single model with dataset as a random intercept, without or with a dataset-specific random SCZ slope; and, set apart at the bottom, the fixed-effect meta-analysis used as the primary analysis presented in the paper in Fig 3 (Methods). Marks give the FDR within each pooled analysis (*** < 0.01, ** < 0.05, * < 0.10, + < 0.20); the number of supertypes at FDR < 0.20 is given at right. The gray row gives between-dataset heterogeneity ($I^2$) from the random-effects meta-analysis. Gray cells, supertype not resolved in that dataset or model not converged. Supertype names are styled as in Fig. 3a (red bold, FDR < 0.10; red italic, FDR < 0.20 in the paper's analysis). (**b**) β under each alternative pooling strategy against β from the paper's fixed-effect meta-analysis, for all 109 supertypes; dashed line, identity; ρ, Spearman; red and labeled, the 11 supertypes at FDR < 0.20 in the paper. (**c**) Leave-one-dataset-out fixed-effect meta-analysis for those 11 supertypes; diamond and bar, estimate and 95% CI from all seven datasets; circles, the seven estimates obtained with one dataset left out.

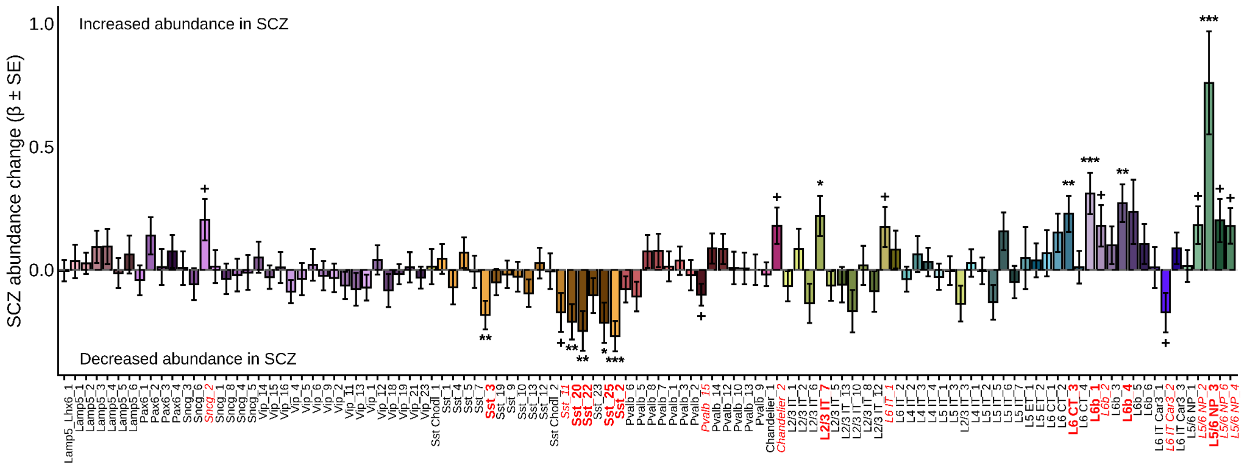

**Fig. S7 | Abundance changes are robust to re-annotation excluding differentially expressed genes.**

Cell type-specific abundance changes in schizophrenia (crumblr β ± SE) after repeating label transfer while excluding genes that were differentially expressed between cases and controls for each of 109 neuronal supertypes meta-analyzed across seven snRNA-seq datasets (N = 298 control / 171 SCZ donors), covarying age, sex and post-mortem interval. Positive bars denote increased cell type abundance in SCZ; negative bars denote decreased or depleted cell types. Bars are colored by supertype. Supertype labels in bold red denote FDR-significant changes (FDR < 0.10) and in italic red trend-level changes (FDR < 0.20); markers above or below bars give *** FDR < 0.01, ** FDR < 0.05, * FDR < 0.10, + FDR 0.10–0.20.

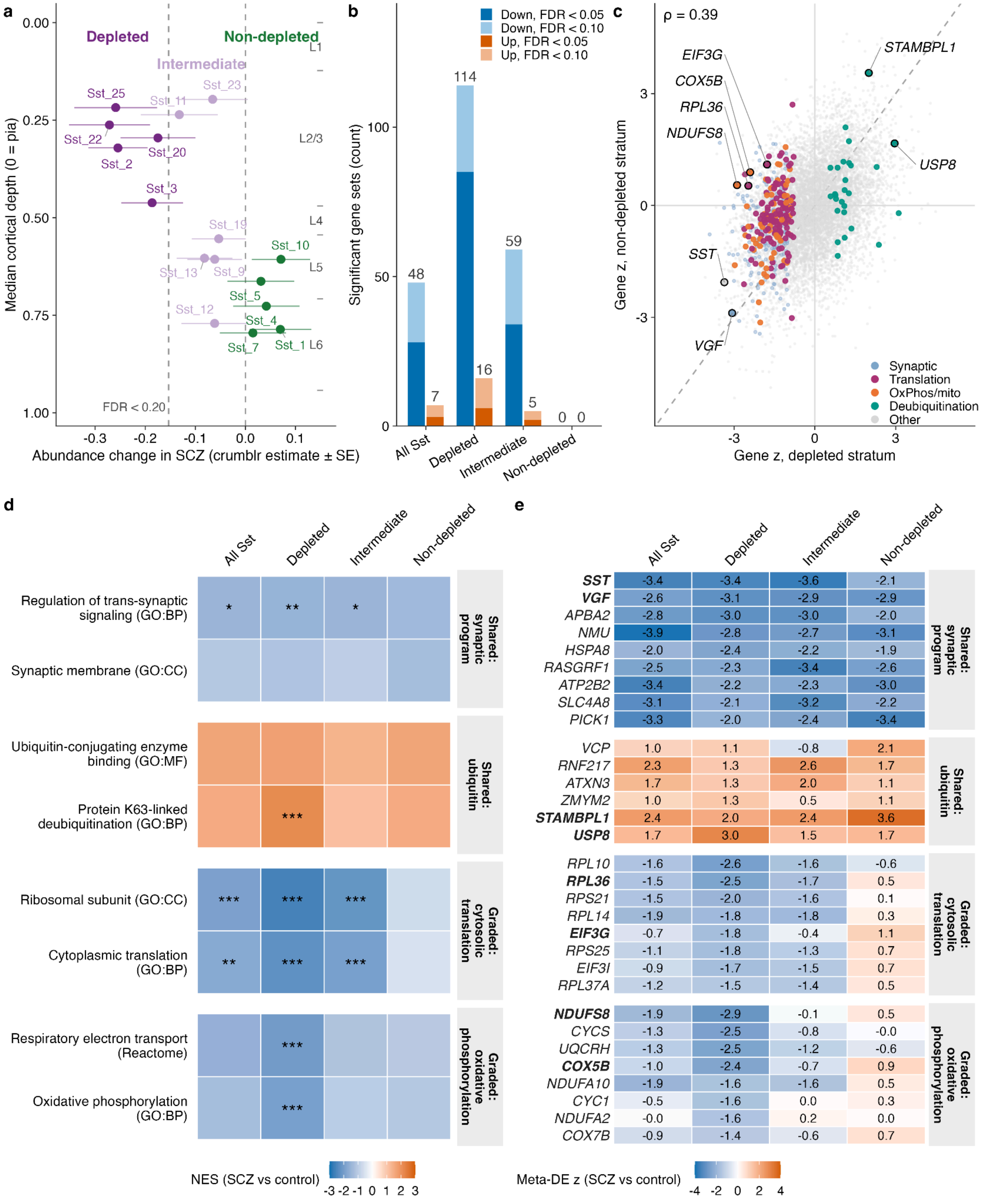


**Fig. S8 | Shared and depletion-graded transcriptional dysregulation of Sst interneurons in schizophrenia.**

(**a**) Definition of the three Sst supertype groups based on depletion status. Per-supertype abundance change in SCZ (crumblr estimate ± SE, from Fig. 3a) against median cortical depth measured in Xenium (0 = pia), for the 16 Sst supertypes. Depleted, FDR < 0.20 (n = 5); intermediate, negative but non-significant (n = 6); non-depleted, estimate ≥ 0 (n = 5). Cortical layer boundaries based on Xenium sections. (**b**) Gene sets significantly enriched in each depletion group; bar labels denote significant gene set counts at FDR < 0.10. "All Sst" denotes the pooled Sst subclass-level analysis of Fig. 2. (**c**) Per-gene meta-analytic z (SCZ versus control) in the depleted versus the non-depleted group, for the shared genes tested in both. Points are colored by membership of the leading-edge union of the gene sets in d; dashed line, identity; ρ, Spearman. labeled genes are shown in bold in e. (**d, e**) Normalized enrichment scores for representative gene sets (**d**) and meta-analytic z for exemplar leading-edge genes (**e**), grouped into blocks shared across the subclass (synaptic, ubiquitin) and blocks graded by depletion (cytosolic translation, oxidative phosphorylation). Significance throughout: ***FDR < 0.01, **FDR < 0.05, *FDR < 0.10.

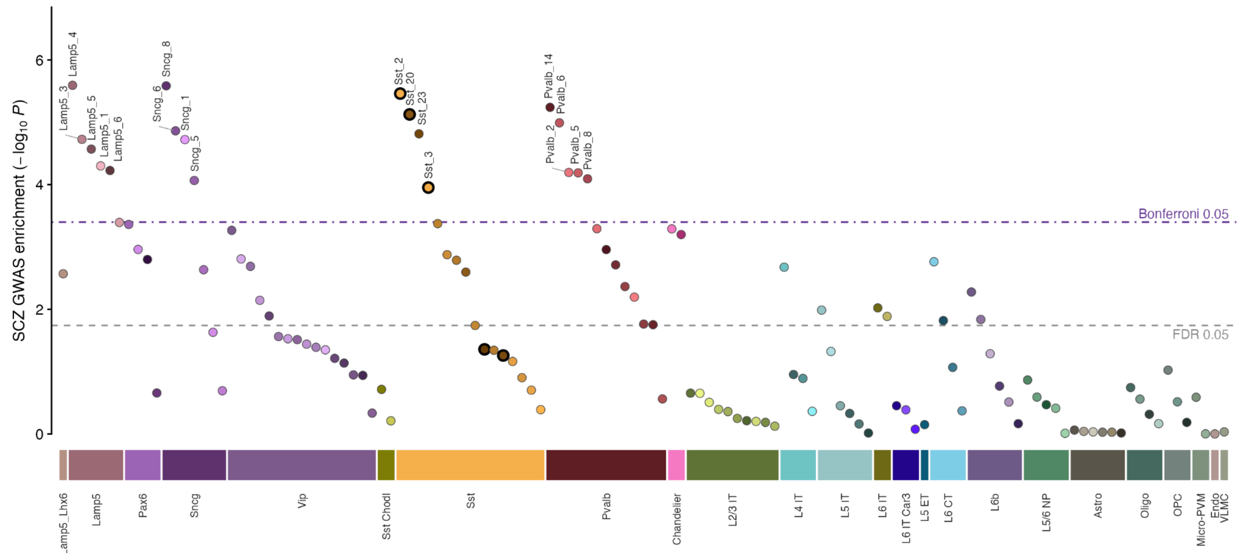


**Fig. S9 | SCZ genetic-risk enrichment across the 125 supertypes of the SEA-AD DLPFC taxonomy.**

The analysis of Fig. 4a extended to every cell type tested: MAGMA gene-property enrichment (one-sided) of SCZ common-variant association (Bigdeli et al. 2026) in the genes specific to each supertype, with expression specificity computed across the 125 SEA-AD DLPFC supertypes (Methods). Each point is one supertype, and all 125 tested supertypes are shown. Supertypes are grouped by subclass (bars beneath the axis, order as in Fig. 3a: GABAergic, then glutamatergic, then non-neuronal) and ordered within subclass by decreasing significance; fill colors are the SEA-AD supertype palette of Fig. 1b. Purple dot-dashed line, Bonferroni P = 0.05 over 125 tests ($-\log_{10}$ P = 3.40); gray dashed line, FDR = 0.05 ($-\log_{10}$ P = 1.74). Supertypes passing Bonferroni are labeled: 18 of 125, all GABAergic interneurons. Of the 51 supertypes at FDR < 0.05, 43 (84%) are GABAergic, eight are glutamatergic and none are non-neuronal. The five Sst supertypes depleted in SCZ (Fig. 3a) are outlined in black; Sst_2, Sst_20 and Sst_3 pass Bonferroni, whereas Sst_25 (P = 0.044) and Sst_22 (P = 0.055) do not reach FDR < 0.05.

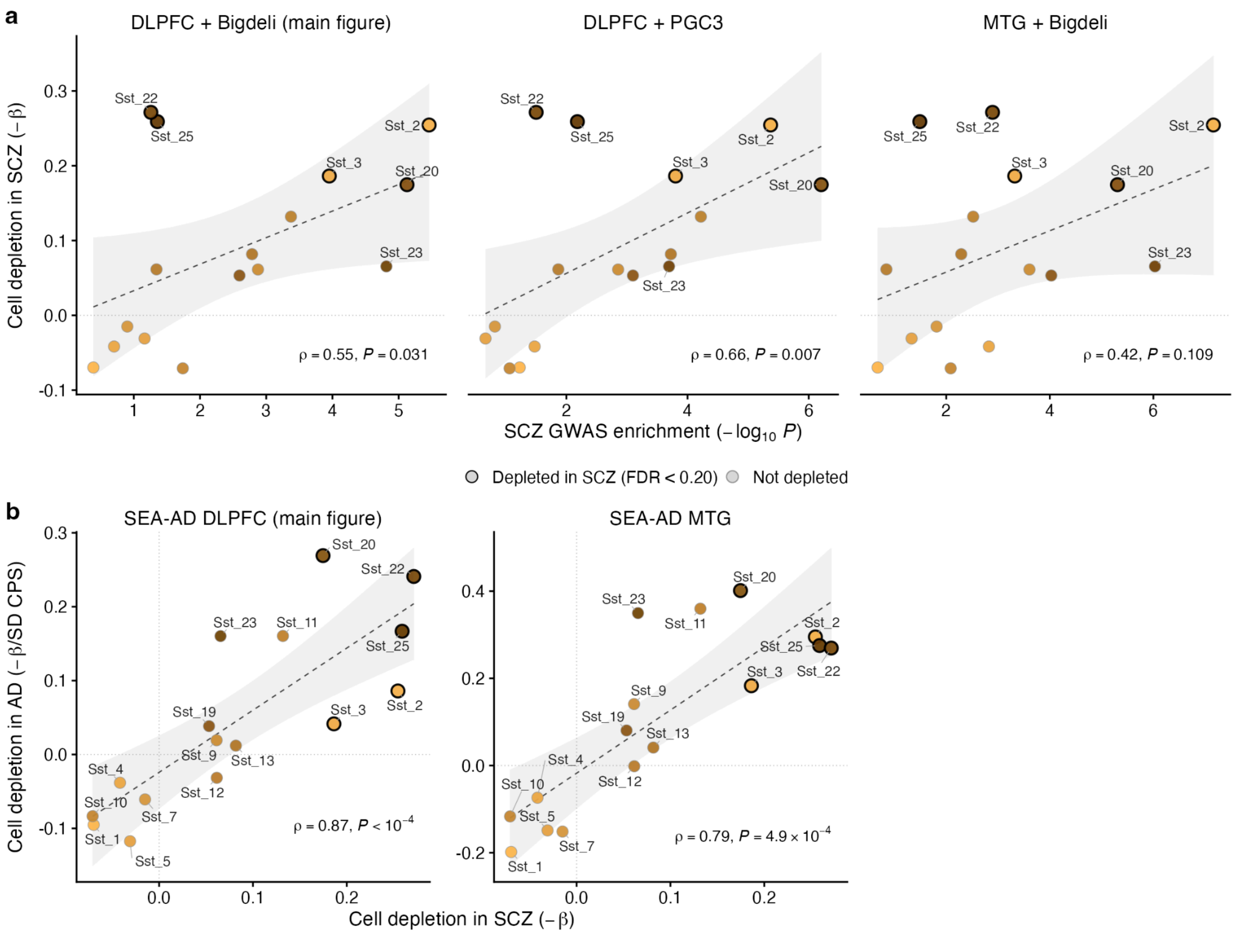


**Fig. S10 | Robustness of the genetic-risk and Alzheimer's disease comparisons to the choice of GWAS and taxonomy reference region.**

Each point is one of the 16 Sst supertypes; thick outlines mark the five depleted in SCZ (FDR < 0.20 in Fig. 3a); dashed lines, linear fits with 95% CI; ρ, Spearman correlation. (**a**) SCZ GWAS enrichment per supertype ($-\log_{10}$ P) versus depletion in SCZ (−β from Fig. 3a), as in Fig. 4a, with the enrichment recomputed substituting each input in turn: the SCZ GWAS (PGC3, Trubetskoy et al. 2022, in place of Bigdeli et al. 2026) and the expression reference (SEA-AD MTG in place of SEA-AD DLPFC, each region contributing its own supertype set — 125 supertypes in DLPFC, 137 in MTG). Specificity calculation, MAGMA settings and compositional data are held fixed. (**b**) Depletion in SCZ (−β) versus decline along the SEA-AD Alzheimer's disease pseudo-progression score (−β per SD of CPS), as in Fig. 4i, computed in DLPFC (main figure) and recomputed in MTG from the same donors with the same model.
