## Supplementary Methods for "Selective depletion of upper-layer somatostatin interneuron subtypes in schizophrenia"

### Xenium cell typing and spatial inference

**Cohort.** We reanalyzed the Xenium dataset of Kwon et al. (2026; GEO GSE307404) comprising 24 post-mortem human DLPFC sections from the LIBD Human Brain and Tissue Repository (12 SCZ, 12 control; matched for age), profiled with a 300-gene panel and yielding 1,339,151 segmented cells. All cell-type, depth, and layer inferences below are analytical additions to this primary dataset.

**Cell-type classification.** With only 300 genes, *de novo* clustering would insufficiently resolve the 24 subclasses and 137 supertypes of the SEA-AD MTG reference (Gabitto et al. 2024), so cell types were assigned using a modified reference-based label transfer approach. We first applied the Allen Institute MapMyCells Cell Type Mapper hierarchical approximate nearest-neighbour (HANN) algorithm (Daniel et al. 2026), a standard tool for mapping cells onto the SEA-AD taxonomy. We then refined the MapMyCells labels using a two-stage classifier built from the Xenium data alone.

Prior to constructing the reference profiles, each Xenium cell's transcript counts were normalized to a common total of 10,000 counts per cell and log-transformed as log(1 + x). For each subclass and each supertype, we averaged expression profiles from the 100 Xenium cells that MapMyCells had assigned to that type most confidently, yielding one reference profile per type. Every Xenium cell was then reassigned to the type whose reference profile the cell most closely matched by Pearson correlation. The reassignment proceeded in two stages: first across the 24 subclasses, choosing the one with highest confidence, and second across only those supertypes belonging to that subclass. Restricting the second stage to within-subclass supertypes enforces consistency between the subclass and supertype assignments, because a cell can never receive a supertype from a different subclass. At the subclass stage we also recorded each cell's correlation with the second-best-matching subclass, and retained the margin (i.e., the difference) between the best and second-best correlation values. We used this as a per-cell measure of classification confidence, that was used subsequently as part of quality control, described below.

*Validation.* Two aspects of the resulting cell type assignments support their validity. First, our subclass calls correspond closely to the original cell type annotations that Kwon et al. generated independently for these same cells (Fig. S2a). For example, our GABAergic subclasses map almost entirely onto their MGE and CGE classes, our excitatory subclasses onto their laminar excitatory classes, and our glial and vascular subclasses onto the corresponding types. Because the Kwon annotations derive from a completely independent clustering of the same cells and dataset, this correspondence is not guaranteed by our procedure. Second, the assigned types express the panel marker genes expected of them (Fig. S2b,d). For example, at subclass level, PVALB, VIP and LAMP5 each reach their highest expression in the corresponding subclass, detected in 92%, 97% and 91% of those cells respectively, while SST marks both Sst and Sst Chodl. At Sst supertype level, Sst_25 is distinguished from the other 15 Sst supertypes by its high expression of CALB1 (detected in 99% of Sst_25 cells), NPY (88%) and NPTX2 (79%). Sst_25 also shows a marked dissociation between the two GAD isoforms, expressing GAD1 in essentially every cell (100%) but GAD2 in only 52%, against 81–97% across the remaining Sst supertypes. This pattern is not an artifact of the Xenium panel or of segmentation, as in the SEA-AD snRNA-seq reference the same Sst_25 supertype expresses GAD1 in 96% of nuclei but GAD2 in only 36%, against 81% across all Sst nuclei.

**Quality control.** Quality control was applied in three steps. First, following Kwon et al., each cell was assessed on transcript-quality and count criteria computed within its own sample. A cell failed if its negative-control-probe, negative-control-codeword, or unassigned-codeword counts exceeded the 99th percentile for that sample; if the number of genes detected fell more than 5 median absolute deviations (MADs) below the sample median; or if its total transcript count fell more than 5 MADs below or above the sample median. Across the 24 sections, 1,298,687 of 1,339,151 segmented cells passed (97.0%; median 97.2% per section, range 93.2–98.2%). Excess transcript counts were the most common failure (23,620 cells), consistent with under-segmented cell pairs.

Second, cells were assessed on the confidence of their subclass assignment. Within each sample, the 5% of cells with the smallest classification margin were flagged. We additionally flagged cells assigned to the L6b subclass whose correlation margin fell below an absolute value of 0.02. L6b is defined by its position at the border of layer 6 and the white matter, yet the initial HANN mapping placed many L6b-labeled cells in upper cortical layers, so we found it empirically helpful to require a greater correlation margin for L6b cells to remove these spatially implausible assignments at the cost of slightly reduced L6b counts. Both thresholds were applied identically to case and control sections.

Third, suspected spatial doublets were flagged by marker co-expression. A cell assigned to a glutamatergic type was flagged if it expressed at least 4 of 7 GABAergic markers (GAD1, GAD2, SLC32A1, SST, PVALB, VIP, LAMP5) and expressed more GABAergic than glutamatergic markers (CUX2, RORB, GRIN2A, THEMIS); a cell assigned to a GABAergic type was flagged if it co-expressed the mutually exclusive markers SST, PVALB, and LAMP5.

Cells passing the margin filter and not flagged as doublets were explicitly marked as being acceptable for downstream analyzes, leaving 1,221,519 cells with a median of 1,068 transcripts and 130 genes detected per neuron, and 527 transcripts and 88 genes per non-neuronal cell.

**Cortical depth.** Each Xenium cell was assigned a normalized cortical depth, running from 0 at the pial surface to 1 at the white-matter border. Rather than estimating depth by manually drawing pial and white-matter boundaries on each of the 24 sections, we predicted depth computationally by transferring it from a reference dataset in which depth had already been annotated by hand. We opted to use the SEA-AD MTG MERFISH atlas to this end (Gabitto et al. 2024), a spatial transcriptomics survey of middle temporal gyrus from 27 aged donors drawn from the larger SEA-AD MTG aged-donor cohort profiled by snRNA-seq. Notably, this MERFISH atlas contains none of the neurotypical reference donors from which the SEA-AD taxonomy itself was derived, so the depth annotations it provides are independent of the reference used to label our Xenium cells. Despite the demographic differences between this dataset and the Kwon Xenium dataset, we selected this MERFISH dataset for two reasons. First, cells in this atlas carry a manually annotated normalized depth from pia. Second, its cells are labeled using the same 24 SEA-AD subclasses and supertypes that were assigned to the Xenium cells, so a cell's local environment can be described in identical terms in both datasets. As cortical layers differ systematically in their cell-type composition, we reasoned that local cell type neighborhood would be informative about the cell's normalized cortical depth; and because the description depends only on the shared subclass labels and not on the gene panel, a relationship learned in MERFISH can be applied directly to Xenium.

Concretely, for each cell we computed the proportion of each of the 24 subclasses among its $K = 50$ nearest neighbours within the same tissue section, and combined those proportions with a one-hot encoding of the cell's own subclass. A gradient-boosting regressor (n_estimators = 300, max_depth = 5, learning_rate = 0.1) was trained on MERFISH cells to map that feature vector onto annotated depth, and the fitted model was then applied to the Xenium cells.

The model was validated by a donor-level split: the three MERFISH donors with the fewest depth-annotated cells were held out for testing and the remaining donors formed the training set, so that performance is measured on individuals the model has never seen. On the held-out donors the model predicted annotated depth with $R^2 = 0.89$ (MAE = 0.069, Pearson $r = 0.95$), against $R^2 = 0.93$ (MAE = 0.050, $r = 0.96$) on the training donors.

Finally, continuous depth was discretized into laminae using boundaries as follows based on visual inspection of the laminar density of excitatory cell subclasses: L1 below 0.12, L2/3 from 0.12 to 0.47, L4 from 0.47 to 0.54, L5 from 0.54 to 0.71, L6 from 0.71 to 0.93, and white matter above 0.93.

**Spatial-domain segmentation.** Cortical laminae were assigned per cell by the depth bins above, but depth alone does not distinguish cortex from adjacent white matter or vasculature. We therefore also partitioned each section into spatially coherent domains using BANKSY ((Singhal et al. 2024; $\lambda = 0.8$, Leiden resolution 0.3, k_geom = 15, 20 principal components), which clusters cells on their own expression together with the expression of their spatial neighbours. Each resulting domain was classified as Cortical, Vascular, or White Matter according to its cell-type composition and its mean predicted depth via the cortical depth classifier defined above. The domains were then used to clean up the per-cell layer calls above: within each domain, isolated layer assignments were replaced by the majority assignment of surrounding cells, removing salt-and-pepper spatial noise. The result partitions every section into cortical laminae L1, L2/3, L4, L5 and L6, plus white-matter and vascular domains.

**Cell types too rare to analyze in Xenium.** We required a cell type to be present in at least half of the donors, which excluded one subclass and four supertypes from our downstream analyzes. The subclass, Lamp5 Lhx6, is reported in our cell-type assignments but was recovered too sparsely in cortex to support per-donor estimates: of the 2,598 cells assigned to it that passed quality control, only 18 lay inside a cortical spatial domain whereas the remainder lay in white matter or vascular annotated domains, and those remaining cortical cells were present in only 11 of 24 donors. Its single supertype, Lamp5_Lhx6_1, is excluded for the same reason. Of the 109 neuronal supertypes carried through the snRNA-seq meta-analysis this leaves 106 supertypes, the others being Pvalb_3, present in 2 donors, and Pvalb_14, which received no Xenium cells at all. Both Pvalb supertypes are poorly resolved by the 300-gene panel (F1 0.42 and 0.49, versus 0.69 and 0.82 using the full transcriptome), so we expect their cells were absorbed into neighboring Pvalb supertypes rather than missing from the tissue.

Among non-neuronal supertypes, the criterion excluded Astro_1 and OPC_, leaving 27 of 29. All exclusions were applied before testing.

**Comparison of cell type assignments against the SEA-AD MERFISH atlas.** We compared our Xenium cell type assignments and predicted depths against the same SEA-AD MTG MERFISH atlas used to train the depth model (341,595 cortical cells, 27 aged donors, 180-gene panel, manual layer annotations). Xenium per-donor subclass proportions agreed with MERFISH at Pearson r = 0.85 ($log_{10}$; ρ = 0.88, n = 23) and median cortical depth at r = 0.96 (subclass, n = 23). Agreement was markedly weaker for the 106 neuronal supertypes (Fig. S3b; r = 0.54, $log_{10}$; ρ = 0.49), reflecting the panel's limited within-subclass marker content. Within each subclass, our model placed different supertypes at systematically different cortical depths, generally consistent with the depths of the same supertypes in the MERFISH data (Fig. S3d,e; median depth r = 0.97 across supertypes and r = 0.94 within subclass). We note however that this depth comparison is not fully independent, since the depth model was trained on the MERFISH annotations from 24 of these 27 donors, but this overall consistency supports that the Xenium cell type assignments generally follow the expected patterns of cortical depths.

**Cell type resolvability given the Xenium panel.** When a supertype (or subclass) fails to show an expected effect in Xenium, either the 300-gene Xenium panel lacks the genes that separate that cell type from its neighbours, or the types are not distinct enough to be separated by any gene set. This distinction bears on how much weight our supertype-level results can carry, and it cannot be settled within Xenium itself: the Xenium supertype labels are the output of the classifier under evaluation, so there is no independent ground truth dataset to score against.

We therefore benchmarked the theoretical performance of the 300-gene Xenium panel directly in the context of the same SEA-AD MTG neurotypical snRNA-seq reference used for cell typing (N = 5 reference donors). We opted to use this dataset for benchmarking because its cell-type labels, including supertypes, are defined from full transcriptomes independently of our work. To this end, we classified nuclei by nearest-centroid Pearson correlation (similar to our Xenium-based classifier described above) under leave-one-donor-out cross-validation and scored the predictions against the reference labels using the F1 score, first on the full transcriptome and then on the 300 panel genes alone. F1 is computed per cell type as the harmonic mean of precision (of the nuclei assigned to a type, the fraction that truly belong to it) and recall (of a type's true nuclei, the fraction recovered), and ranges from 0 to 1, where 1 denotes perfect recovery of that type with no misassignments. Here, holding the classifier, cross-validation scheme, and reference labels fixed leaves the gene set as the only variable to be tested.

We found that subclass identity was generally very well predicted using the 300-gene Xenium panel in this benchmark (median F1 0.96 with the Xenium panel vs 0.99 with the full transcriptome; 15 of 23 subclasses at ceiling performance; Fig. S2c). However, Sst supertype identity was only partly resolved using the 300-gene Xenium panel versus the full snRNA-seq-based transcriptome (median F1 0.62 vs 0.88; Fig. S2e). The high transcriptome ceiling shows the Sst supertypes are genuinely separable given enough genes, so the gap in F1 scores reflects Xenium panel coverage rather than biological continuity. We reason that this is because the panel carries very few within-subclass discriminating markers for most Sst supertypes (Fig. S2d). Nevertheless, Sst_25 was the best-resolved (F1 0.81), whereas Sst_22

(F1 0.45) and Sst_20 (F1 0.44) were the worst-resolved, potentially accounting for the non-replication of the Sst_22 and Sst_20 composition effects in Xenium.

We note that this snRNA-seq-based benchmark is not a direct measurement of Xenium performance, as snRNA-seq nuclei are not subject to segmentation errors, spatial doublets, or the potentially reduced per-cell detection sensitivity of imaging-based data; thus these F1 values are an upper bound on what the panel can achieve in Xenium rather than the accuracy we attain in practice. Given these limits on cell typing at supertype resolution with a restricted gene panel, we treat analyzes making use of the Xenium supertype cell type calls as corroborative of the snRNA-seq findings rather than definitive.
